# Convergent stromal and immune remodeling defines spatial tumor dynamics in PARP inhibitor–resistant high-grade serous ovarian cancer

**DOI:** 10.64898/2026.03.03.709035

**Authors:** Kathleen Jane Imbach, Sergi Cervilla Garcia, Daniela Grases, Sara Bystrup, Arola Fortian, Adrià Bernat-Peguera, Mustafa Sibai, Lorena Valdivieso Almeida, Elvira Carballas, Pau Guillén Sentís, Margarita Romeo, Eduard Porta Pardo, Jordi Barretina

**Author notes:** These authors contributed equally (Kathleen Jane Imbach, Sergi Cervilla Garcia).

## Abstract

Acquired resistance to poly(ADP-ribose) polymerase inhibitors (PARPi) remains a major barrier to durable clinical benefit in patients with high-grade serous ovarian cancer (HGSOC), yet the profiles of tumors at resistance remain poorly understood. We profiled longitudinal HGSOC patient tumors spanning diagnosis, post-neoadjuvant therapy, and progression on PARPi with complementary spatial transcriptomics platforms, integrating single-cell–resolution Xenium analysis in paired longitudinal cases with cohort-level Visium profiling. PARPi failure was associated with spatial remodeling of the tumor microenvironment, marked by hypoxia-associated malignant transcriptional programs, increased stromal compartmentalization, and exclusion of effector immune cells. Together, these findings indicate that PARPi resistance in HGSOC is accompanied by reproducible spatial niche remodeling, underscoring the tumor microenvironment as a key contextual determinant of therapeutic failure.

## Introduction

Ovarian Cancer (OC) is the leading cause of gynecological cancer-related deaths in women, with a survival rate of less than 50% after five years.^1,2^ Most OC patients are diagnosed with the aggressive high grade serous (HGSOC) subtype characterized by high degree of molecular heterogeneity and near ubiquitous loss of functional TP53, with 20% of patients harboring BRCA1/2 mutations and almost half exhibiting homologous recombination deficiency (HRD).^3^ The standard first-line treatment for patients with advanced HGSOC consists of systemic platinum and paclitaxel-based treatment, with surgical debulking when feasible.

Significant advances have been made with the introduction of poly (ADP-ribose) polymerase (PARP) inhibitors (PARPi) as maintenance therapy, primarily in HRD patients.^4–6^ Despite positive initial responses to therapy, most patients ultimately show disease progression due to the emergence of resistance. Known mechanisms of PARPi resistance, including restoration of homologous recombination function via secondary mutations in BRCA1/2 or in other HR-related genes,^7^ along with other alterations in DNA repair pathways, explain only part of this clinical failure. This suggests that resistance may not be fully accounted for by cancer cell genetics, but may also reflect constraints imposed by the broader tumor ecosystem.

Numerous studies have explored HGSOC heterogeneity using bulk and single-cell RNA sequencing approaches, revealing transcriptional diversity among malignant, immune, and stromal populations.^3,8–11^ However, these methods rely on dissociated tissue and do not explore the tumor architecture. The development of spatial transcriptomics (ST) has permitted profiling of tumors in the intact context of the tumor microenvironment (TME), helping to reveal how spatial patterns contribute to cancer progression, immune evasion, and therapeutic resistance. Initial applications of ST in HGSOC have characterized distinct molecular clusters linked to chemotherapy response,^12,13^ as well specific tumor-immune cell interactions.^14–17^ However, these studies focus on treatment-naive or post-neoadjuvant chemotherapy (NACT) samples, even when trying to associate data with PARPi response.^17^ As a consequence, the spatial organization of tumors following PARPi progression remains poorly characterized.

Different spatial platforms provide complementary insights; single-cell–resolution technologies enable detailed analysis of local architecture and cellular communication, while transcriptome-wide methods permit cohort-level estimation of genomic events and pathway activity. Here, we investigate how HGSOC tumors are reorganized at progression on PARPi by applying a multi-platform spatial transcriptomics strategy. Integrating single-cell–resolution Xenium analysis in longitudinal sample pairs with cohort-level Visium profiling, we test the hypothesis that therapeutic resistance follows conserved spatial constraints even when malignant evolution is heterogeneous. By characterizing how cancer cell populations, stromal remodeling, and immune cell positioning converge within intact human tissue, we elucidate spatial principles that contextualize resistance and shape tumor evolution in HGSOC.

## Results

To characterize spatial remodeling associated with PARPi resistance, we assembled a longitudinal cohort of HGSOC samples from 11 patients obtained at: diagnosis during primary debulking surgery (n=5), after NACT at interval debulking surgery (n=5), or at secondary cytoreductive surgery upon progression on PARPi (n=6). We captured complementary characteristics of tumor organization with multi-platform ST approaches; Xenium afforded single-cell–resolution analysis in selected longitudinal cases, while Visium permitted transcriptome-wide profiling across the cohort (**Fig. 1a**). Samples were obtained from ovary, omentum, peritoneum, and lymph node sites, and comprised paired longitudinal cases and additional unpaired samples spanning treatment states (**Fig. 1b**). Detailed clinical data for each patient, including BRCA status and initial platinum therapy response, are detailed in **Table S1**.

**Figure 1.**
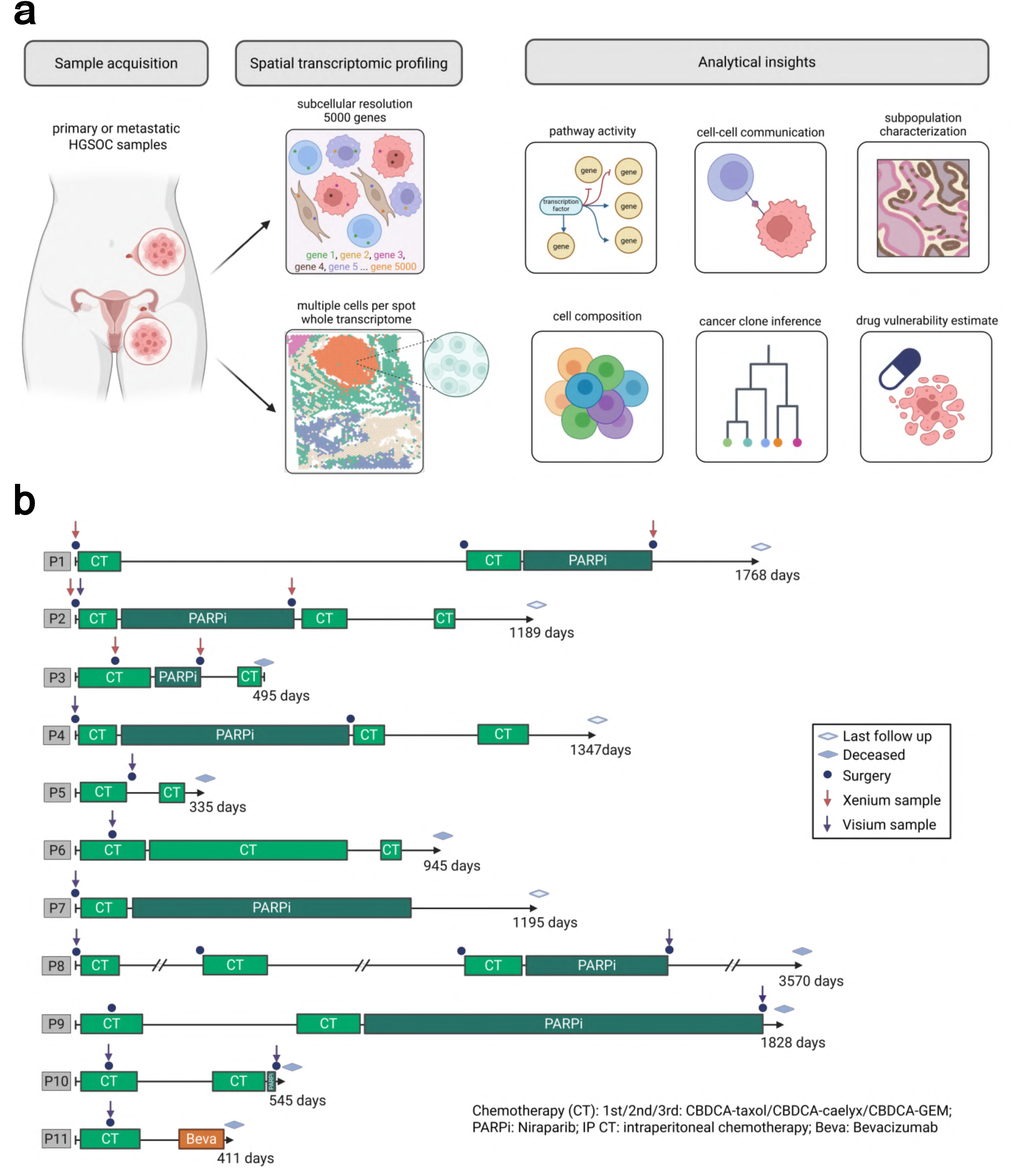
A. Multi-platform spatial profiling strategy. Complementary spatial technologies balance resolution and breadth; 10x Genomics Xenium Prime 5K resolves single-cell spatial architectures, subcellular transcript localization, and microenvironmental niches, while 10x Genomics Visium captures whole-transcriptome spatial domains, copy number variant (CNV) profiles, and drug sensitivity signatures. B. Clinical cohort overview. Treatment and sampling timelines of the 11 HGSOC patients (P1-P11), with arrows emphasizing at which surgical resection the samples were collected for spatial transcriptomics profiling, colored according to the platform utilized. CT: Chemotherapy; PARPi: PARP inhibitor therapy; Beva: Bevacizumab.

### Resistant tumors feature composition and neighborhood changes

We first examined how tumor composition differed between pre- and postPARPi samples using the high-resolution ST data. While treatment-naive tumors exhibited cohesive nests of cancer cells forming papillary or micropapillary structures, the NACT-treated patient (P3) showed a more intermingled distribution with stromal and immune cells **(Fig. 2a).** PARPi-treated tumors exhibited a similar shift toward fragmented, dispersed patterns, supported by the overall relative decrease in malignant contribution at this timepoint **(Fig. 2b)**.

**Figure 2.**
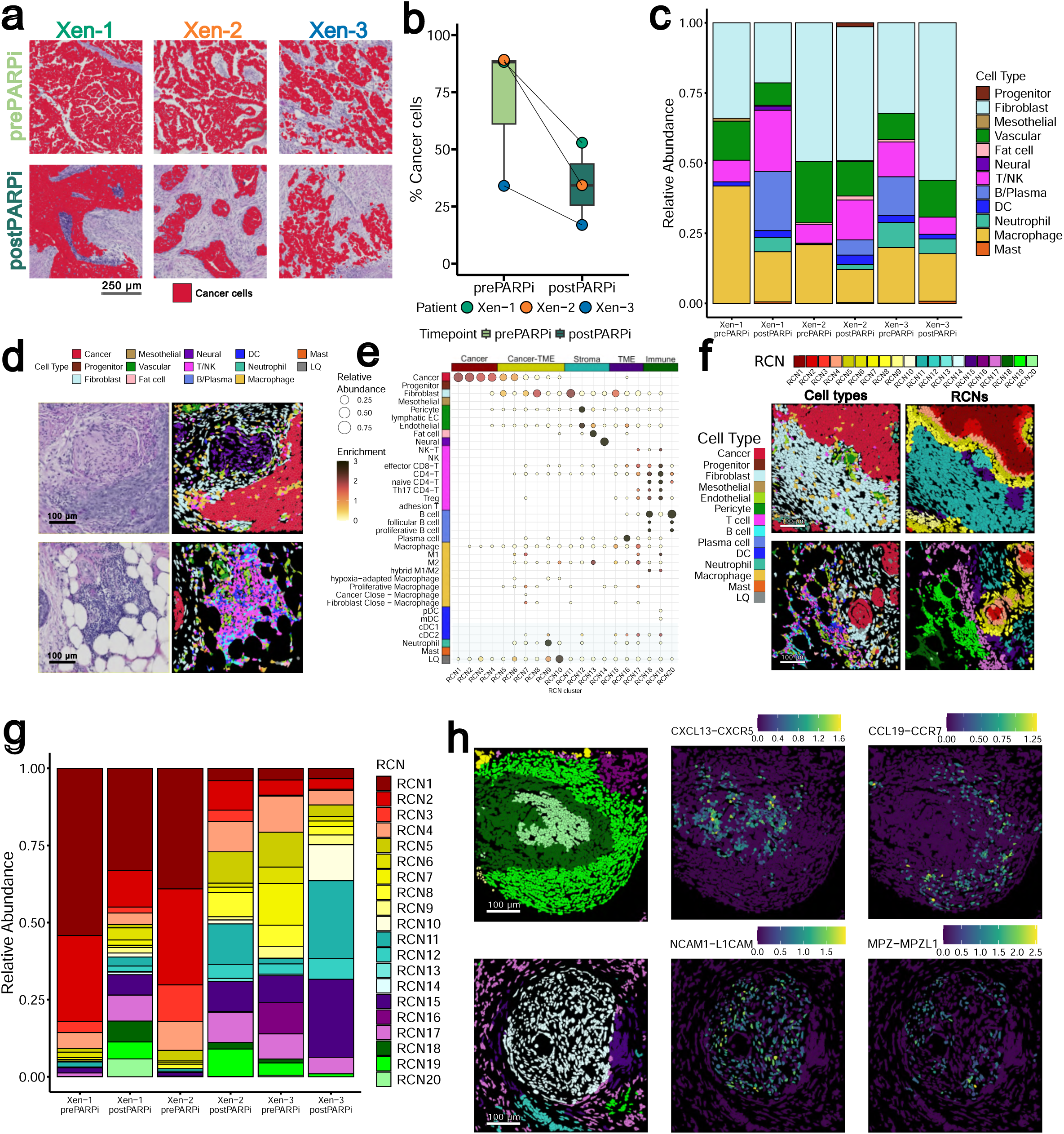
A. Representative H&E images with segmented cells classified as malignant overlaid in red for each sample profiled with Xenium, with patients in columns and timepoints in rows. B. Boxplot showing the percent of Xenium samples composed of malignant cells by timepoint. Lines drawn connect corresponding timepoints for each patient. C. For the non-malignant compartment, stacked bars show the relative abundance of each cell type per sample. D. Examples showing cell segmentation areas colored by the cell identity assigned to each alongside the corresponding H&E area for cancer-proximal peripheral nerve bundle in Xen-1 postPARPi (top) and fat-associated tertiary lymphoid structure in Xen-2 postPARPi (bottom). E. A bubble plot shows the granular cell contributions to each RCN 1-20. Bubbles are sized according to the relative proportion of contribution of that cell within the RCN context and colored according to enrichment. F. Tissue examples show cell segmentation colored by cell type (left) and corresponding RCN assignments (right) for Xen-1 postPARPi (top) and Xen-2 postPARPi (bottom). G. A stacked bar plot shows the relative abundance of cellular RCN assignments within each of the samples. H. Inferred LR signaling in two RCN contexts: CXCL13-CXCR5 and CCL19-CCR7 in TLS RCNs 18-20 in Xen3 postPARPi (top); NCAM1-L1CAM and MPZ-MPZL1 in RCN 14 in Xen1 postPARPi.

TME cell characterization highlighted the pronounced heterogeneity of the tumors **(Figs. 2c and S2a-S2b),** with a few trends observed between treatment states. Treated samples showed an increase in general immune cell contribution, with neutrophils appearing only in NACT or PARPi treated samples and mast cells present only in post-PARPi samples (**Fig. S2c**). Evaluation of annotated cells in space revealed structures previously described in HGSOC, including tertiary lymphoid structures,^18^ peritumoral nerve formations,^19^ and cancer cell invasion into the mesothelial layer^20^ **(Fig. 2d)**.

We subsequently used local cellular composition and spatial proximity to capture patterns of higher-order spatial organization, defined here as recurrent cellular neighborhoods (RCNs; Methods). The 20 total RCNs grouped tissues into tumor cores (RCNs 1-4), tumor-TME interfaces (RCNs 5-10), and stroma- or immune-enriched regions (RCNs 11-20; **Fig. 2e-f**). Comparing RCN distributions across treatment states corroborated treated samples (NACT and PARPi) to exhibit loss of cancer-specific structures but an increase in immune structures (**Fig. 2g**), suggesting that PARPi progression is associated with reproducible changes in spatial architecture.

We then assessed the functional biological relevance of the RCNs obtained by evaluating signaling patterns therein. We evaluated estimated intercellular ligand-receptor (LR) interactions within these spatial contexts to quantify domain-specific differences in cellular cross-talk (**Fig. S2d**; Methods). Notable examples included *CXCL13-CXCR5* and *CCL19-CCR7* in tertiary lymphoid structures (TLS) captured by RCNs 18-20,^21–23^ and *NCAM1-L1CAM* and *MPZ-MPZL1* in peripheral nerve RCN 14 (**Fig. 2h**).^24,25^

Together, these analyses establish that HGSOC tumors undergo global spatial remodeling after PARPi failure, manifested in tissue composition changes and neighborhood organization. This descriptive framework provides the foundation for subsequent analyses aimed at identifying the microenvironmental interactions and biological programs underlying these spatial changes.

### Malignant profiles highlight spatially-clustered transcriptional heterogeneity and hypoxia-associated tumor niches after PARPi progression

Having established that PARPi progression is accompanied by tumor configuration shifts, we next focused on malignant cells to determine cancer-intrinsic features of the longitudinal tumors. We first assessed the global malignant transcriptomic patterns across samples by aggregating Xenium transcriptomes by sample and running principal component (PC) and differential expression (DE) analyses (Methods). PC space stratified samples by initial therapy and patient, but less so by timepoint **(Fig. 3a)**. Notably, within-patient transcriptomic distances in the PC space correlated with progression-free survival (PFS), underscoring the temporal dimension to intratumoral evolution. DE analysis of bulked expression by PARPi status yielded 78 post-PARPi and 85 pre-PARPi differentially expressed genes (DEGs) without significant pathway enrichment; however, notable genes such as *STAT3* and *SLC7A5*, were significantly upregulated after PARPi, in line with previous results^26,27^ **(Table S2)**.

**Figure 3.**
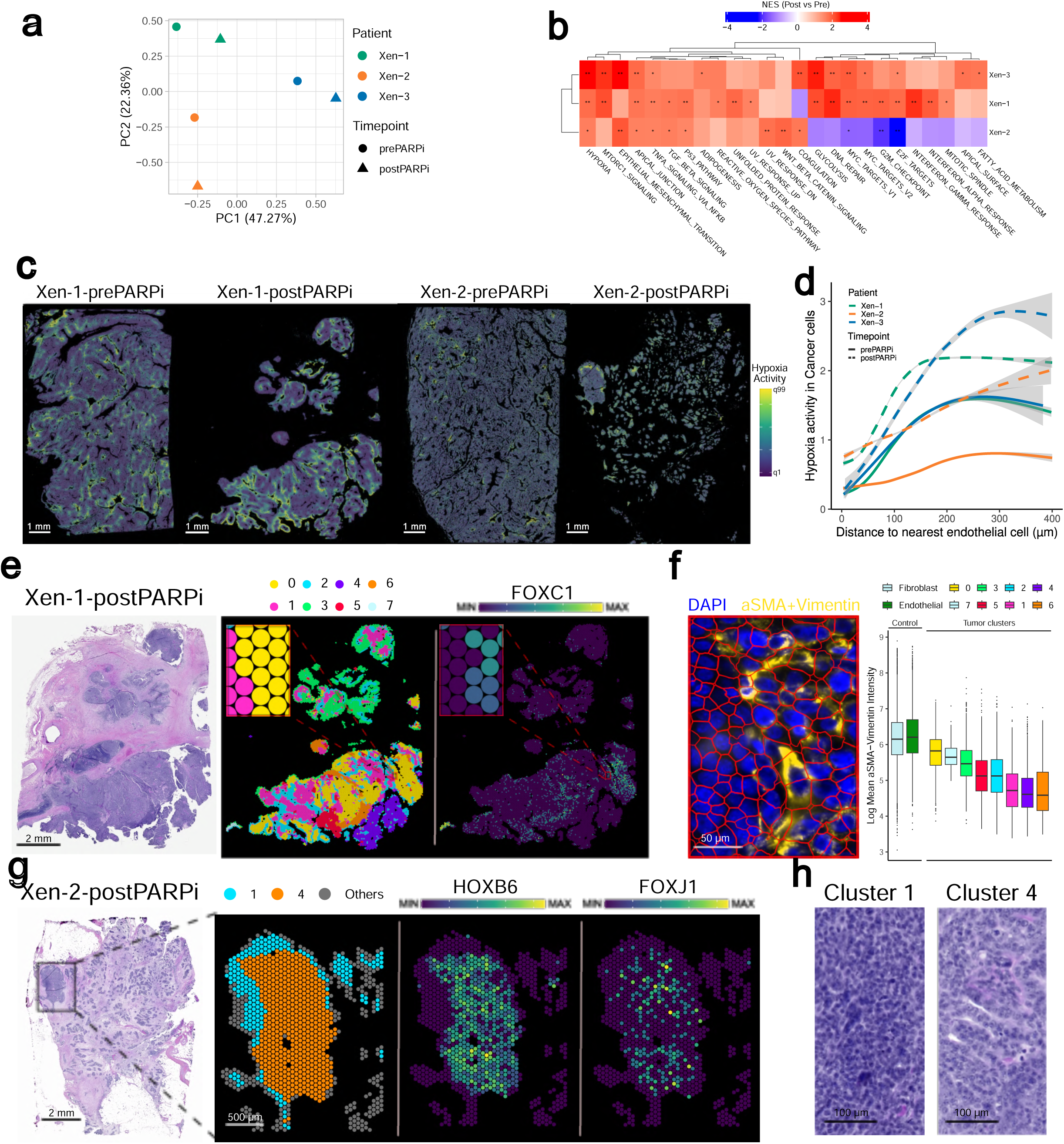
A. Principal component (PC) plot for the pseudobulked expression of malignant cells per sample, with points colored by patient and shaped by timepoint. B. Heatmap showing the normalized enrichment score (NES) for relevant MSigDB Hallmarks after conducting differential expression per-sample between post-PARPi and pre-PARPi bulked malignant gene profiles. FDR-adjusted p-values are shown where significant (*: adjusted p-value < 0.05; **: adjusted p-value < 0.01). C. Progeny hypoxia pathway scores overlaid on the malignant areas of each tissue, with segmented malignant cells colored by estimated pathway activity (scale ranges from the first quantile, q1, to the 99th quantile, q99). D. The relationship between the hypoxia activity scores in cancer cells and their corresponding distance to the nearest endothelial cell in µm. Lines are colored by patient and stroked by timepoint. Gray areas show confidence intervals. E. The H&E image of the full sample area for Xen-1-postPARPi (left) alongside its corresponding malignant rasterization, with raster pixels colored by their assigned cluster identity (middle) and their expression of *FOXC1* (right) with the same area highlighted in the inset panels of each. F. Left: the same area as the inset panel shown in (H) for Xen-1-postPARPi, this time with the DAPI (blue) and aSMA+Vimentin (yellow) protein staining shown and the cell segmentation overlaid in red. Right: boxplots show the distribution of log10 mean aSMA-Vimentin staining intensity within segmented cells either of fibroblasts and endothelial cells or malignant cells assigned to each of the rasterized clusters from Xen-1-postPARPi. G. Left to right: the H&E image of the full sample area for Xen-2-postPARPi, a zoomed area of the largest cancer cell block divided into malignant raster pixels and colored by raster clusters (clusters 1 and 4 highlighted), expression of *HOXB6*, and expression of *FOXJ1*. H. Zoomed H&E examples of Xen-2-postPARPi areas where malignant clusters 1 and 4 were defined highlight differences in cell morphology.

We then asked how established cancer-associated pathways might differ between treatment timepoints. To address this, we compared longitudinal changes within each patient and scored MSigDB Hallmark pathways^28^ (Methods). *Hypoxia*, *apical junction*, and *TNF*α *signaling via NF-*κ*B* were significantly up-regulated in all three patients following PARPi (**Fig. 3b**). In contrast, P2 displayed down-regulation of *G2M checkpoint*, *E2F targets*, and *MYC targets*, suggesting a less proliferative and more quiescent state. The idiosyncrasy of P2 persisted when evaluating canonical HGSOC markers in individual cells (**Fig. S3a**); this patient’s post-PARPi sample exhibited relatively lower expression of HGSOC-specific marker genes (*MUC1*, *MUC16*, *PAX8*, and *SOX17*), but was unique in its expression of neuronal related genes (*CADM3*, *SLITRK4*, *TUBB3*, *NEFH*, *CHRNA3*, *CHRNB4*) (**Fig. S3a**). This contrasted strongly with P3, whose malignant cells expressed high levels of genes associated with cell proliferation and cycling (*MKI67*, *BIRC5*, *TOP2A*, *UBE2C*).

Having established overall differences in cancer activity by timepoint, we next probed how cancer programs vary on the cellular level. For this, we leveraged the PROGENy pathways^29^ for inference. Comparing sample distributions of the malignant cells’ pathway scores further supported a consistent hypoxia-associated signal in post-PARPi tumors (**Fig. S3b**), with the extent of this difference more pronounced in the samples with lower PFS (**Fig. S3c**). Mapping the hypoxia scores onto tissue sections revealed localized areas of enriched values (**Fig. 3c**), leading us to ask whether the hypoxic signature might be related to oxygen gradients imposed by intratumoral vasculature patterns. When comparing the hypoxia score of each cancer cell with its distance to the nearest endothelial cell, we found that cancer cells further from endothelial cells present higher hypoxia levels, especially beyond 150 µm (**Fig. 3d**). This gradient was accentuated in post-PARPi samples, with cancer cells exhibiting sharper increases and larger absolute hypoxia scores at the same distances as their primary counterparts.

Finally, we sought to profile inter- and intra-tumor phenotypic heterogeneity within the cancer cells in a data-driven manner. We applied a spatial pseudobulking strategy by aggregating malignant cells within a 55 µm diameter to create spatially conserved transcriptional programs (Methods). These spatial aggregates were then used to determine malignant-specific clusters per sample. Several pseudobulk-derived clusters shared common profiles, such as those exhibiting interferon (IFN) related signaling and others defined by proliferation/cell cycling genes (**Fig. S3d**). Other clusters demonstrated distinctly patient- and region-specific characteristics. In P1 post-PARPi, we identified spatially restricted malignant clusters enriched for epithelial–mesenchymal transition (EMT)–associated programs, including *FOXC1* and *TGFB3*, localized to defined tumor regions (**Fig. 3e**). The clusters were concordant with increased Vimentin and αSMA protein expression quantified for Xenium-based cell segmentation, supporting the mesenchymal-like phenotype of these spatially-defined malignant cells (**Fig. 3f**).

We also observed a spatially-confined subpopulation of tumor cells exhibiting lineage plasticity in P2 post-PARPi (**Fig. 3g**). Located in a single large tumor nest, the cluster overexpressed genes associated with developmental and epithelial remodeling, including *HOX* family members (*HOXB6*, *HOXB5*, *HOX9*, *GATA4*, *PAX2*), and multiciliation (*FOXC1*, *CCNO*, *DNAH5*). This cluster was spatially segregated from an adjacent proliferative malignant population expressing canonical cell-cycle genes (*CCNA2*, *TK1*, *E2F1*). Transcriptional divergence was accompanied by overt histological differences, as corresponding H&E morphology revealed a more structured, differentiated architecture in the *HOX*-high region compared with densely packed proliferative tumor (**Fig. 3h**).

Together these findings underscore the heterogeneity within the malignant compartment of HGSOC from a transcriptional and morphological standpoint. Cancer cells of P3 were characterized by gene signatures related to proliferation, while spatially-constrained subgroups of post-PARPi cancer cells from P1 and P2 exhibited evidence of EMT and lineage plasticity, respectively. Finally, we observed that hypoxia pathway activity consistently increased following PARPi therapy, even when accounting for proximity to vasculature. Together, these results highlight the wide range of phenotypes that can be adopted by malignant cells in an individual lesion, while also pointing to hallmark cancer pathways that consistently define post-PARPi trends.

### Signatures of CAF activation and the stroma’s role as a physical barrier

In addition to the malignant cells themselves, fibroblasts represent a major component of the HGSOC TME and have been implicated in shaping cellular interactions and tissue patterns in the progression of the disease.^30,31^ To understand how PARPi progression may impact stroma remodeling, we scrutinized fibroblast states and their spatial organization between treatment timepoints.

Analysis of fibroblast expression bulked by sample revealed consistent transcriptional shifts following PARPi progression. In PC space, the stromal signatures showed strong division by PARPi status and tissue (**Fig. 4a**). DE analysis revealed that post-PARPi fibroblasts adopted a cancer-associated fibroblasts (CAF)-like transcriptional program characterized by increased expression of extracellular matrix genes (*ASPN*, *COMP*, *ELN*, *TNXB*, *COL4A4*), together with activation of mechanotransduction and contractile signaling (*MRTFA*) and pro-fibrotic Hedgehog and STAT3 pathways (*GLI1/GLI2*, *PTCH1*, *STAT3*).^32–34^ In contrast, primary timepoint fibroblasts were more heterogeneous but exhibited immune modulation genes (*IL10, IL4, IL13, LIF, CSF3*), consistent with a more immunoregulatory rather than myofibroblastic stromal state. Notably, expression of the bulk-implicated genes was heterogeneous across individual fibroblasts, consistent with activation of specific fibroblast sub-states rather than a uniform transcriptional response in the tissue stroma (**Fig. 4b**).

**Figure 4.**
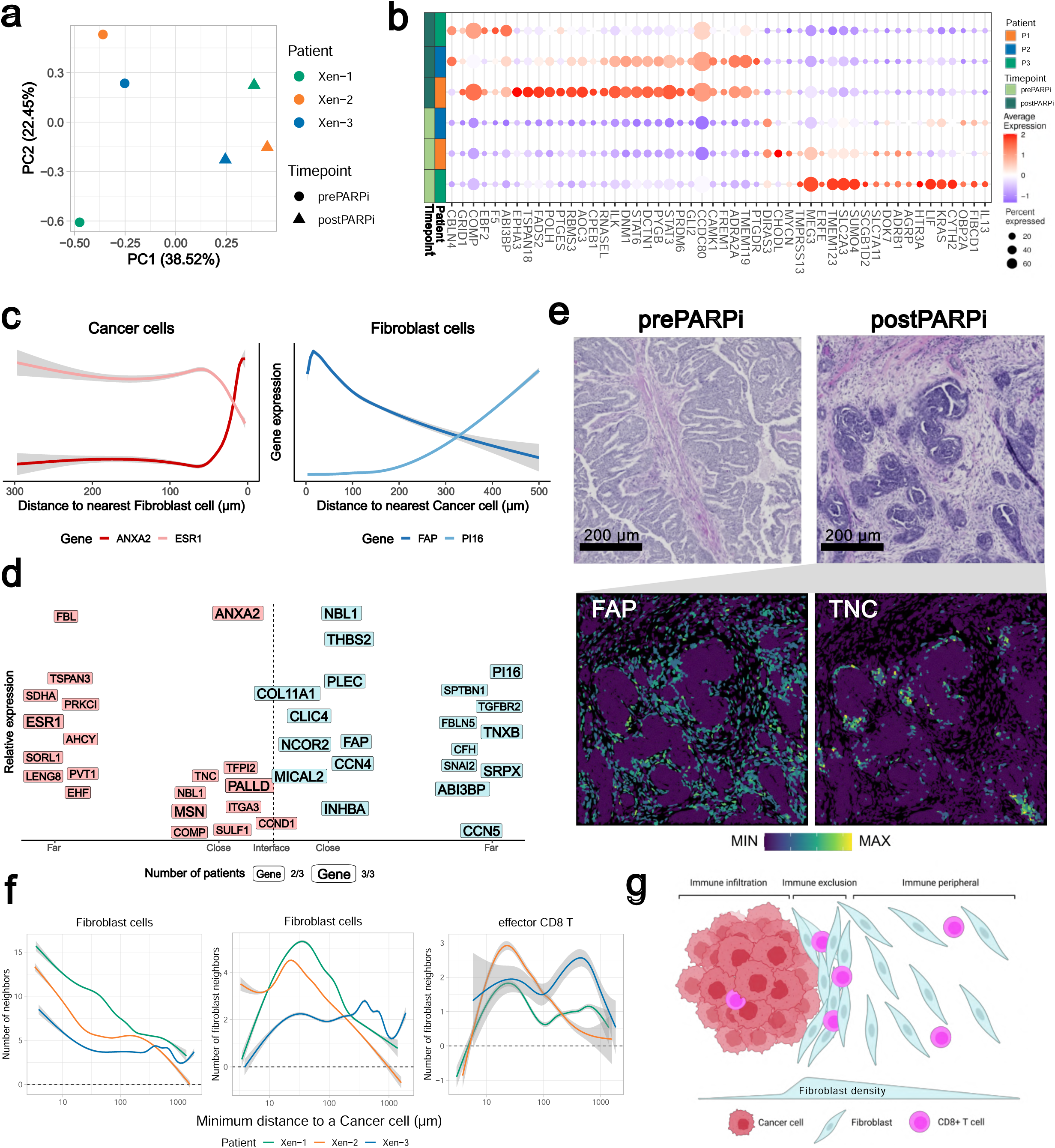
A. Principal component (PC) plot for the pseudobulked expression of fibroblast cells per sample, with points colored by patient and shaped by timepoint. B. A bubble plot highlights the top differentially expressed genes in fibroblasts between timepoints for each sample, with dot size representing the percent of total fibroblasts expressing the gene and the color denoting the average expression in fibroblasts. C. Line graphs show the change in gene expression of ANXA2 and ESR1 within cancer cells as they get nearer to fibroblasts (left), and conversely the expression of FAP and PI16 in fibroblasts in relation to distance to cancer cells (right). Lines are colored by genes and confidence intervals are shown in gray for each line. D. The most consistently observed genes that change with location on the cancer-stroma interface axis in the post-PARPi samples, with cancer-specific genes shown on the left in red and fibroblast-specific genes shown on the right in blue. Genes are positioned on the y-axis according to their relative expression, and genes with a larger typeface correspond to those observed in all 3 samples while those with a smaller typeface are observed in 2. E. H&E of tissue areas showing the morphologies of fibroblasts near malignant cells for Xen-1 at each timepoint (top), with the post-PARPi timepoint area overlaid with cell segmentation colored by POSTN expression (bottom left) and estimated TNC-ITGB6 activity (bottom right). F. Line plots highlight the relationship between minimum distance to a cancer cell and the total number of neighbors in fibroblasts (left), the number of fibroblast neighbors in fibroblasts (middle) and the number of fibroblast neighbors in CD8+ T cells (right) within a 25µm radius. Lines are colored by patient (post-PARPi only) and the gray regions show confidence intervals. G. A cartoon depicts a working model of the stroma as a physical barrier, with dense fibroblasts surrounding the tumor, thereby excluding CD8 T cells.

To spatially contextualize the treatment-associated differences in fibroblasts, we modeled gene expression as a function of proximity to malignant regions in the post-PARPi samples, distinguishing tumor cores, tumor–stroma interfaces, and distal stromal areas (Methods). The proximity-based analysis revealed distinct spatial gradients in fibroblast and cancer gene expression, with specific expression programs preferentially enriched at the tumor–stroma interface compared with both tumor cores and distal stroma (**Fig. 4c-e, Table S3a**). Notably, interface-associated fibroblast genes tended to recur across samples, whereas those on the cancer side were more sample-specific (**Fig. 4d**).

Extending the proximity analysis beyond individual genes to LR signals in all samples revealed fewer interactions shared consistently across patients (**Table S3b**). In cancer cells near fibroblasts, only *ITGA3–LAMB3*/*LAMC2* interactions were recurrent across post-PARPi samples. In tumor-adjacent stroma, *POSTN* interactions with integrins (*ITGB3/5, ITGAV*) and *THBS2–ITGAV* were shared across all post-PARPi samples and the P2 pre-PARPi sample. Distal fibroblasts showed more variable interactions, including *CXCL12–ACKR3* in two of three post-PARPi samples and *THBS1–CD36* in all samples except P3 post-PARPi, highlighting greater heterogeneity in distal stromal signaling.

The transcriptional-based observations were supported by histological evidence, with post-PARPi samples exhibiting increased stromal density and desmoplastic architecture surrounding tumor nests compared to pre-treatment samples (**Fig. 4e**). Dense cancer-associated stroma has been proposed to act as both a physical and functional barrier to immune cell infiltration.^35^ To assess stromal organization quantitatively, we examined the relationship between cellular neighborhood density within a 25 μm radius and distance from cancer cells (**Fig. 4f, Fig. S4**; Methods). Cancer-adjacent fibroblasts tended to have higher overall neighbor density, while fibroblast–fibroblast neighbor counts peaked at approximately 30 μm from tumor cells. Effector CD8L T cells similarly exhibited a peak in fibroblast neighbor density at approximately 25 μm from cancer cells, suggesting that immune cell positioning is constrained within regions of elevated stromal density (**Fig. 4g**).

### Progressive HGSOC features immune exclusion, spatially constrained inflammation hotspots, and *CXCL12-CXCR4* signaling

Given the consistent stromal remodeling observed following PARPi progression, including increased stromal density, we next examined how the immune landscape is altered in these tumors. Although immune cells have been implicated in tumor progression and therapeutic resistance in HGSOC,^36^ these tumors are typically “immune-cold” and patients experience limited clinical benefit from immunotherapy.^37,38^ We therefore assessed how immune infiltration and organization change following PARPi progression.

Immune cells and structures increased in treated samples (**Fig. 2c**), but their presence alone doesn’t indicate effective proximity to malignant regions. We first quantified T cell infiltration in the samples, comparing the number of T cells found in tumor-rich RCNs compared to the total number of non-TLS T cells present (Methods), and found that T cells are increasingly excluded from tumor cores in post-PARPi timepoints (**Fig. 5a**). Then, we leveraged the intercellular geometries to probe where key effector populations, CD8+ T cells and M1 macrophages, are found in relation to malignant cells in the tissues. Post-PARPi samples displayed a redistribution of these immune cells away from malignant regions (**Fig. 5b**), underscoring the importance of studying tumors within their spatial contexts. These results suggest that immune exclusion may be exacerbated in HGSOC following PARPi administration, despite the relative increase in immune cell contribution to the TME following therapy.

**Figure 5.**
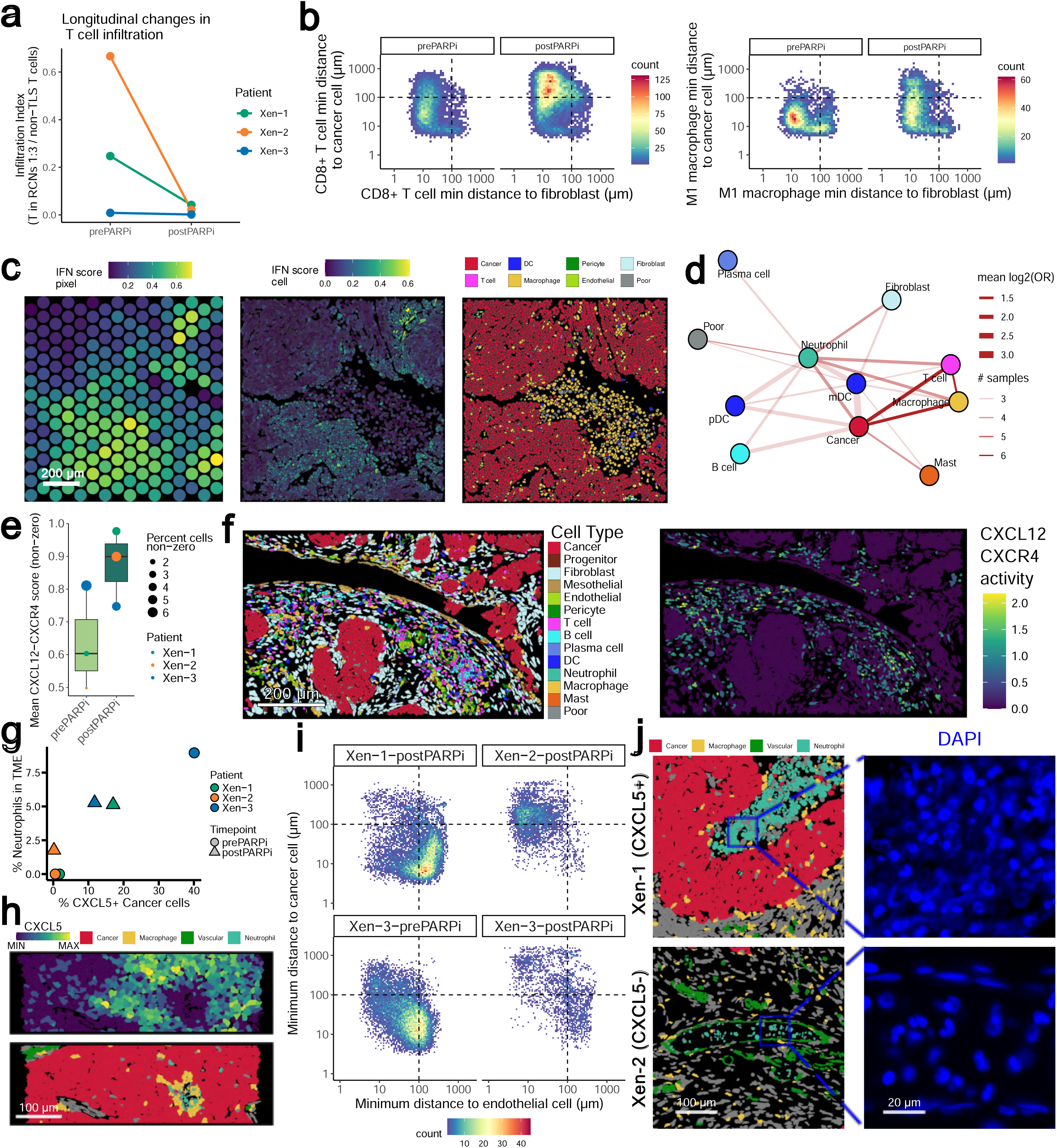
A. A dot plot shows the infiltration index, calculated by the proportion of T cells found in RCNs 1-3 relative to the total number of T cells not found in TLS RCNs, at each timepoint. Dots and lines connecting samples are colored by patient. B. Density plots show the number of CD8+ T cells (left) and M1 macrophages (right) and located relative to cancer and fibroblast cells at pre-PARPi and post-PARPi timepoints. C. An example tissue area from Xen-1-prePARPi showing the IFN pathway activity on the rasterized pixels (left), in the individual cells (middle) and the corresponding cell type annotations (right). D. A network plot highlights the co-localization of cell types in areas with high IFN activity in the samples, with cell types represented as nodes and the mean log2 odds ratio (log2OR) of a co-occurrence shown by line opacity and the line thickness denotes the number of samples in which the occurrence was observed. E. A boxplot shows the mean CXCL12-CXCR4 scores in cells with appreciable levels by timepoint. Dots are sized according to the proportion of cells within that sample with non-zero scores and are colored by patient. F. An example tissue area from Xen-1-postPARPi showing segmented cells colored by annotation (left) and estimated CXCL12-CXCR4 activity in each cell (right). G. A scatterplot shows the relationship between the proportion of cancer cells expressing CXCL5 and the percent contribution of neutrophils to the total TME, with points shaped by timepoint and colored by patient. H. Tissue area of Xen-1-postPARPi showing the cell segmentation colored by CXCL5 expression (top) and cell annotation (bottom). I. Density plots show the number of neutrophils found relative to the nearest endothelial and cancer cells in the samples where neutrophils were detected. J. Left: tissue areas with cell annotation for Xen-1 (top) and Xen-2 (bottom) show neutrophils near cancer cells and within a blood vessel, respectively. Right: DAPI staining of zoomed areas from regions in the left panels.

Despite the evidence of immune exclusion, we had previously observed cancer subsets recurrently expressing IFN-associated genes across samples (**Fig. S3d**). We asked how these regions of inflammation were distributed in the tissue, and which cells tended to occur there (**Fig. 5c**, Methods). Mapping IFN response activity revealed that inflammatory signatures were confined to discrete regions often on the periphery of malignant regions, decoupled from large tumor cores. In IFN-high regions, cancer consistently coincided with macrophage and T cells, while neutrophils, mast cells, and mDCs were also enriched in select samples (**Fig. 5d**). Together, these patterns point to restricted regions of inflammation hotspots rather than distributed immune activation in these tumors.

Finally, given the presence of the *CXCL12-CXCR4* chemokine axis in RCNs 16-19 and its reported relevance in OC cells (**Fig. S2d**),^39–41^ we probed how this signal might differ between timepoints. Both the proportion of cells with appreciable *CXCL12-CXCR4* activity and the average activity scores themselves were more pronounced in treated samples, including the pre-PARPi NACT-treated case of P3 (**Fig. 5e**). Evaluation of the cells contributing to these scores per sample particularly pointed to immune cells (**Fig. S5a**), and mapping the scores onto the tissue highlighted the activity in the immune and stroma cells adjacent to malignant cell clusters (**Fig. 5f**). These results suggest that the *CXCL12-CXCR4* signaling axis may be an additional characteristic of the surrounding TME in resistant tumors.

### Neutrophil heterogeneity and spatial distribution suggest NETosis as a feature of treated samples

After observing that effector immune cells are present within the tissue but spatially displaced from malignant niches after PARPi treatment, we next examined neutrophils in greater depth. Neutrophils were detected exclusively in NACT- and PARPi-treated tumors (**Fig. 2c**, **Fig. 5g**), suggesting a potential association with disease progression or therapeutic response.

Spatially, neutrophil abundance in the TME correlated with expression of the neutrophil-recruiting chemokine *CXCL5*^42,43^ on malignant cells (**Fig. 5g-h**). As an exception, in patient P2—who exhibited the longest PFS—*CXCL5* expression was absent in cancer cells despite neutrophil presence, indicating the presence of *CXCL5*-independent recruitment mechanisms.

Spatial analysis revealed two major neutrophil localization patterns: intravascular neutrophils and neutrophils in direct contact with cancer cells (**Fig. 5i**). Tumor-proximal neutrophils preferentially expressed genes associated with inflammatory and hypoxic responses (***IL1B*, *IFI16*, *HIF1A*, *HK2*, *CTS***) whereas tumor-distal or intravascular neutrophils expressed genes related to neutrophil activation (***FCGR3A/B*, *CSF3R*, *SELL***; **Table S4a)**. The specific transcriptional patterns defining these spatially distinct states varied between patients, indicating heterogeneous spatial neutrophil states instead of a uniform gene signature.

Consistent with the transcriptional differences, the distinct neutrophil states exhibited variation in nuclear morphology by location and *CXCL5* status. In *CXCL5*-expressing tumors, tumor-proximal neutrophils exhibited diffuse, decondensed DAPI staining, whereas intravascular neutrophils in *CXCL5*-negative tumors displayed compact, lobulated nuclei (**Fig. 5j**). This chromatin decondensation pattern is consistent with DNA scaffolds observed in neutrophil extracellular traps (NETs), which function primarily to trap and neutralize pathogens in a process termed NETosis.^44,45^ Although transcripts associated with NETs^46,45^ (*MPO*, *ELANE*, *PADI4*) were not detected (**Fig. S5b**), neutrophils from patients P1 and P3 displayed increased nucleus-based segmentation relative to membrane-based segmentation, consistent with chromatin expansion and poorly defined cellular boundaries characteristic of NETs^46,45^ (**Fig. S5c**).

Neutrophil co-localization with other immune and stromal cell types varied across patients; however, post-PARPi samples from patients P1 and P3 displayed highly similar interaction patterns (**Fig. S5d**). In both cases, neutrophils co-localized preferentially with hypoxia-adapted macrophages and conventional dendritic cell subsets (cDC1 and cDC2). Notably, neutrophils exhibiting NET-like morphological features were enriched within hypoxic regions, which also contained low-quality (LQ) cells—potentially reflecting cells undergoing NETosis, characterized by chromatin extrusion and RNA degradation. The observed co-localization patterns were consistent with spatial niches RCN9–10.

Finally, we examined whether the proximity of neutrophils influenced transcriptional programs in neighboring cancer cells. In addition to the upregulation of hypoxia-associated genes, consistent with the localization of NET-like neutrophils in hypoxic niches, neutrophil-adjacent cancer cells exhibited patient-specific inflammatory and stress-related signatures. In P1, neutrophil proximity was associated with expression of *CCL20*, *SERPINE3*, and *SDC4*, whereas in P3, proximity corresponded to expression of *CCL28*, *ANGPTL4*, and *HK2 (***Table S4b).**

Together, these findings demonstrate that neutrophils occupy spatially and transcriptionally distinct niches within the TME and reinforce the relevance of NETs as a potential immunomodulatory and pathological feature in HGSOC.

### Tissue remodeling patterns are reinforced across spatial modalities

While the Xenium data enabled detailed insights into spatial tumor remodeling associated with PARPi at single-cell resolution, it profiled a limited set of samples and covered a restricted subset of the transcriptome. To examine cohort-level patterns of tumor organization, we leveraged complementary Visium spatial transcriptomics. Although inter-patient heterogeneity remained substantial (**Fig. 6a**), Visium profiled samples recapitulated treatment associated changes in tissue composition observed previously.

**Figure 6.**
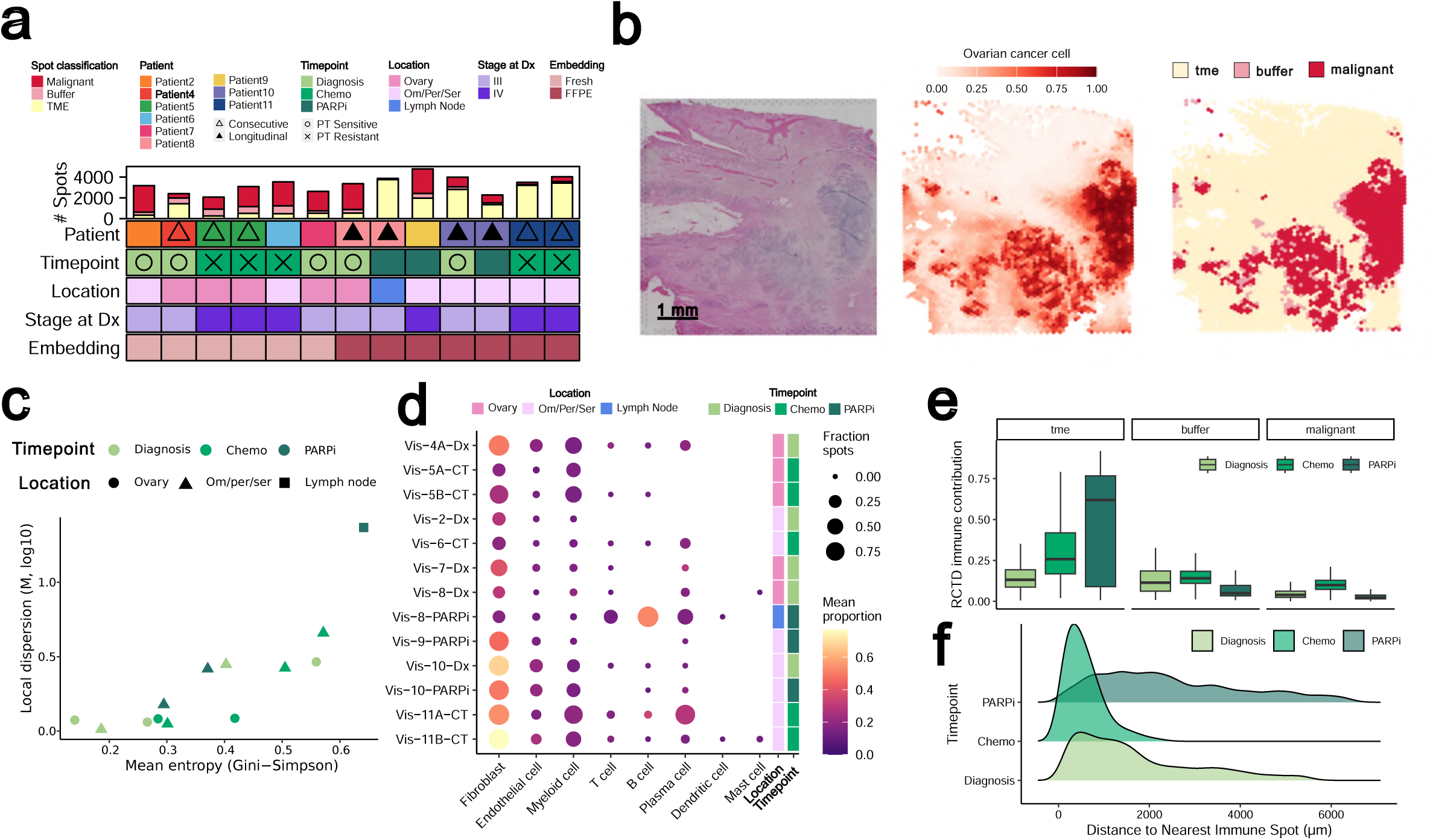
A. A heatmap shows the sample attributes of those tissue sections profiled with Visium, with a stacked bar plot atop denoting the total number of Visium spots assigned to each compartment. Chemo: chemotherapy treated (NACT or otherwise); Om/Per/Ser: omentum/peritoneum/serosa; Dx: diagnosis; FFPE: formalin-fixed paraffin-embedded. PT: platinum therapy. B. Tissue example for Vis-8-Dx showing the H&E (left), the per-spot estimation of ovarian cancer cell contribution (middle), and the final spot classifications (right). C. Scatterplot depicts the mean entropy of deconvolution scores (Gini-Simpson) on the x-axis versus the log-transformed local dispersion (Marcon and Puech’s M, radius of 400µm) for the malignant spots per Visium sample. Points are colored by timepoint and shaped by tissue location. D. A bubble plot showing the deconvolution scores of TME spots per Visium sample, with dot size corresponding to the fraction of TME spots with non-zero estimations and the color denoting the mean proportion in non-zero spots. Tiles to the right are colored by tissue location (left) and timepoint (right). E. Boxplots showing the summed immune estimates per spot in each of the compartments, with boxes separated and colored by sample timepoint. F. A ridge plot of the distribution of distance of the nearest immune spot (summed proportion > 0.25) in µm to each malignant spot, stratified by sample timepoint.

First, cellular contributions per spot were estimated with both reference-based and reference-free approaches (Methods, **Fig. 6b**). Classification into TME, buffer (mixed malignant and TME), and malignant compartments using the reference-based estimates were validated with the reference-free estimates (**Fig. S6a**). We subsequently evaluated the structural trends of the malignant regions in each tumor by assessing the entropy, or extent of “mixing”, within the malignant spots and how dispersed those spots were in the broader tissue context (Methods). Across samples, treated cases tended to exhibit higher levels of intraspot mixing and less spatial concentration in the tissue (**Fig. 6c**), mimicking the fragmented structure of cancer cells in treated samples observed with Xenium. In the non-malignant compartment, scrutiny of cell contributions in TME by sample showed that location strongly influenced contributions; lymphocyte signatures predominated in a sample obtained from a lymph node while fibroblasts and myeloid cells were the main estimated components of all the other samples (**Fig. 6d**).

Given the Xenium-backed evidence that treated samples tend to have a higher contribution of immune cells, we evaluated the aggregated immune estimates within each spot across samples (Methods). When stratifying spot immune proportions by compartment and treatment timepoint, we similarly observed that chemotherapy (CT) and PARPi-treated samples have higher immune estimates in their TME. However, post-PARPi samples in particular lacked immune signatures in their buffer or malignant zones (**Fig. 6e**). To further interrogate the spatial configuration of the immune compartment in the tumors, we calculated the proximity between immune-high spots with malignant spots in each sample (Methods). Cancer spots in samples taken after CT exhibited closest proximity to immune regions, while the immune distance was skewed to be further from malignant areas in diagnosis and post-PARPi samples (**Fig. 6f**).

Though of a lower spatial resolution, the Visium data reiterated the shifts in tumor configurations observed at single-cell; CT- and PARPi-treated tumors tend to have higher contributions of immune cells in the TME than treatment-naive samples, but samples after PARPi treatment in particular feature immune signatures distally located from tumor regions and lack evidence of infiltration.

### Whole-transcriptome data permits estimation of malignant clones and drug sensitivity predictions, providing clinically relevant insights

After recapitulating the PARPi-associated tumor remodeling patterns from a compositional perspective, we asked how we might utilize the Visium data to glean additional insights beyond those possible with Xenium’s limited gene set. Using the full transcriptome, we focused on the malignant compartments to more deeply profile pathway activity, infer cancer clones, and evaluate signatures of drug sensitivity.

Previous work has demonstrated that spot-level ST signatures can be used to estimate large scale copy number variants (CNV) in malignant tissues.^47^ When this approach was applied in HGSOC samples, high confidence CNVs correlated well with corresponding whole genome sequencing data (WGS).^15^ Using the expression profiles from malignant spots, we inferred clones in each sample by comparing the malignant area to a published healthy ovarian ST dataset^48^ (Methods).

The number of unique clones per sample ranged from one to 8, and did not correlate with the number of malignant spots in a sample (**Fig. S7a**). Scrutiny of the most commonly affected driver genes in samples highlighted established HGSOC-associated events, such as amplification of *MYC* and loss of *RB1* (**Fig. S7b**).^3,49^ Similarities of CNV landscapes between each possible pair of clones demonstrated that clones from the same patient tended to be more similar, even between timepoints (**Fig. 7a**). Comparing inferred genome-wide events in two pairs of samples (Methods), one adjacent pair taken at the same timepoint and a longitudinal case from a PARPi-treated patient, further supported this observation; while the CNV profiles of the adjacent pair’s clones intermixed, those of the longitudinal pair showed clear separation by timepoint, with the single PARPi progression clone serving as an outgroup from all diagnosis timepoint clones **(Fig. S7c**).

**Fig 7.**
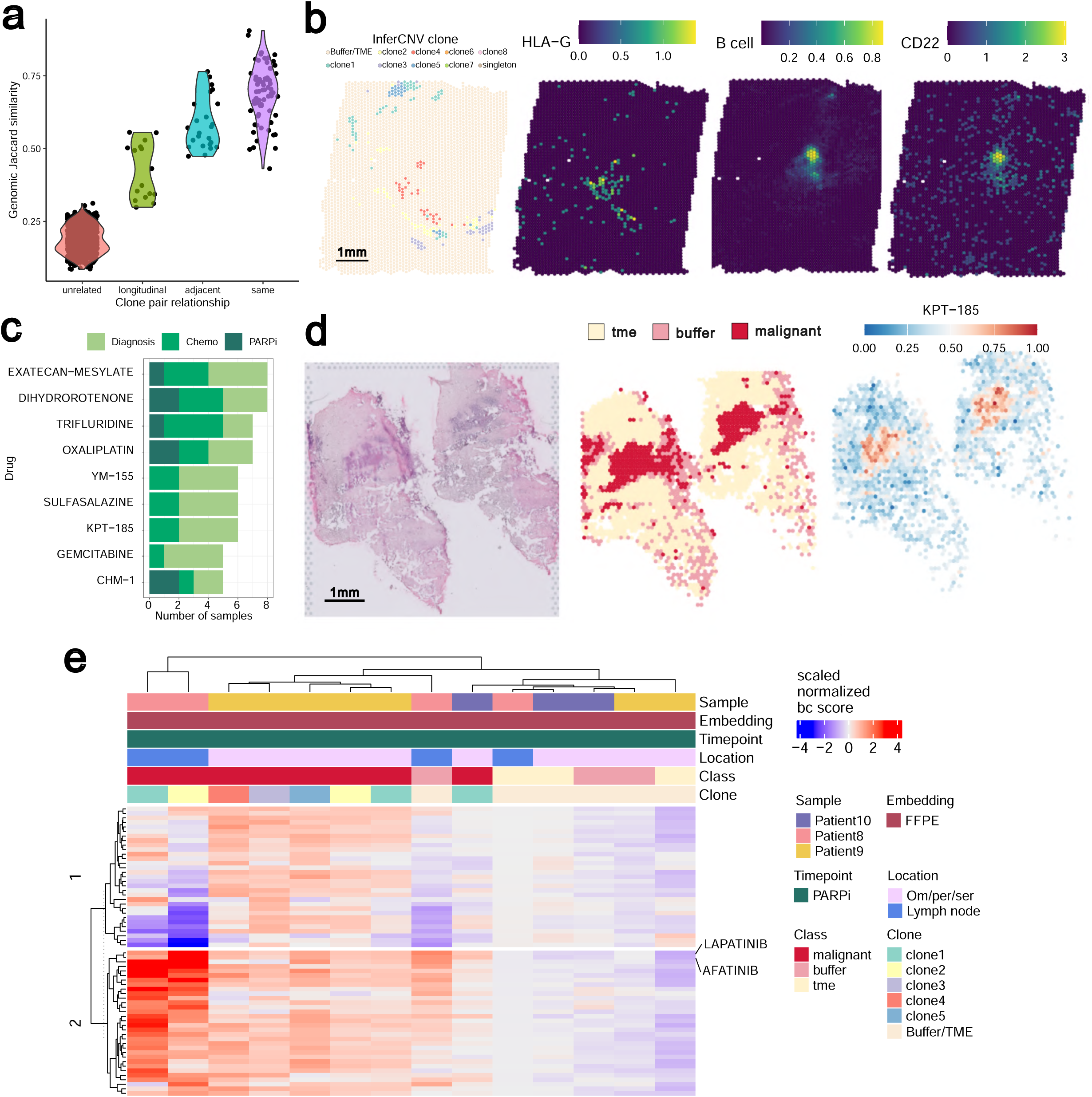
A. Violin plot showing the Jaccard similarity between clone pairs’ estimated CNV profiles according to the relationship between the samples. B. Tissue overlay plots of Vis-9-CT-A showing, from left to right: InferCNV-derived clone annotations, the expression of *HLA-G*; B cell contribution per spot, the expression of *CD22*. C. A stacked bar chart shows the top BeyondCell drug signatures predicted for the samples, with the x-axis denoting the total number of samples implicated, the y-axis indicating the drug name, and the bars colored according to the treatment timepoint of the samples. D. Tissue plots of Vis-2-Dx corresponding to sample H&E (left), the RCTD-based spot annotations (middle), and the predicted sensitivity to KPT-185 per spot (right, spots with NA values resulting from the model are not shown). E. A heatmap of all the BeyondCell drug signature scores for all signatures predicted as selectively targeting malignant areas of post-PARPi samples, split by clonal compartments. Signature scores are split into 2 blocks by clustering, with the rows corresponding to Lapatinib and Afatinib highlighted.

Not unlike the malignant subclustering approach in the Xenium, grouping the Visium tissues by clone and profiling gene expression enrichment permitted detection of subpopulation patterns that would have been otherwise missed. Notably, clone-specific differential expression analysis revealed that an inferred clone from Vis-11A-CT was highly enriched for expression of *HLA-G* (average log2 fold change > 4.96, FDR < 3.1e-81; **Fig. 7b; Table S5**), a gene encoding an immunosuppressive class I histocompatibility antigen involved in fetal tolerance^50^ and reported to be overexpressed in tumors.^51^ Proximal to these clones in the TME was a region enriched for B cell composition and expression of *CD22*, an inhibitory receptor responsible for curbing B cell reactivity^52^ (**Fig. 7b**).

Finally, we used the full transcriptome data to predict drug vulnerabilities in the Visium samples using BeyondCell^53^ (Methods). Classical cytotoxic drugs, such as anti-microtubule drugs, DNA damaging agents, and replication inhibitors scored as top drug candidates in the samples (**Fig. 7c**). Interestingly, drugs in clinical trials for ovarian cancer were also implicated. For example, the nucleocytoplasmic transport inhibitor KPT-185 which selectively inhibits Exportin 1 showed high sensitivity score in the malignant areas of 4 naïve and 2 interval samples.

Though limited by the number of samples, we asked which drug dependencies were predicted in PARPi-resistant tumors in particular. In two of the 3 post-PARPi samples profiled with Visium, BeyondCell predicted sensitivity to ErbB inhibitors Lapatinib (HER2i) and Afatinib (pan HERi) (**Fig. 7d**). Unlike other predicted drugs, the scores were consistently elevated across all malignant clones in the post-PARPi samples (**Fig. 7e**).

## Discussion

In this study, we examined how standard therapies reshape the spatial organization of HGSOC, with a focus on architectural and ecosystem-level features. By leveraging complementary spatial transcriptomics platforms across longitudinal patient samples, our work demonstrates how therapy alters tumor composition, cellular interactions, and microenvironmental structure in ways that would be difficult to infer from bulk or dissociated single-cell approaches alone.

Treated tumors consistently exhibited disruption of cohesive papillary tumor nests accompanied by increased prominence of stromal and immune compartments. Importantly, while both datasets indicated increased immune cell presence following therapy, spatial context revealed that PARPi-resistant tumors displayed segregation between malignant and immune compartments. This observation highlights a critical limitation of non-spatial profiling approaches: immune cell abundance alone may misleadingly suggest enhanced immunogenicity,^54^ whereas spatial context reveals pronounced immune exclusion. This distinction has direct clinical relevance, as multiple trials have evaluated combinations of PARP inhibitors and immunotherapies in relapsed HGSOC with limited benefit,^55–57^ underscoring that immune cell abundance alone does not reflect effective anti-tumor engagement.

When we interrogated local architectures and cell-specific transcriptional states, we found that malignant transcriptional programs remained highly patient-specific, consistent with prior reports in HGSOC.^58,59^ These findings reinforce the challenge of identifying universal cancer-intrinsic vulnerabilities in this disease. Notably, one post-PARPi sample exhibited a spatially restricted malignant population characterized by expression of developmental regulators and epithelial remodeling genes alongside reduced expression of canonical HGSOC markers. This phenotype is consistent with lineage plasticity or partial de- or trans-differentiation and has important translational implications. As antibody–drug conjugates and other targeted therapies increasingly rely on lineage-restricted antigens,^60^ spatially localized loss of these markers may constitute a mechanism of therapeutic escape and would likely be diluted or undetectable in bulk RNA analyses.

Among the most consistent malignant-associated shifts observed following PARPi exposure was the induction of hypoxia-related transcriptional programs. Hypoxia estimates were not uniformly elevated, but intensified with increasing distance from endothelial cells, a pattern that was more pronounced in post-PARPi samples. This spatial relationship suggests that therapy-associated hypoxia may arise from the emergence of newly formed, poorly vascularized tumor regions rather than global changes in oxygen demand. Given established links between hypoxia, genomic instability, immune suppression, and therapy resistance,^61–64^ reinforcement of hypoxic niches following PARPi may contribute to both malignant persistence and broader microenvironmental reprogramming.

In contrast to the pronounced heterogeneity of malignant programs, stromal remodeling emerged as a more conserved feature of treated tumors. Fibroblasts in post-PARPi samples upregulated gene expression programs consistent with CAF-like states,^33,34,32^ and spatial analyses revealed that transcriptional programs at the tumor–stroma interface were more consistent on the stromal side than within adjacent malignant compartments. This asymmetry suggests that while malignant states evolve idiosyncratically, stromal responses may represent a more conserved and potentially targetable consequence of therapy. Histological and spatial analyses further supported this interpretation, revealing denser, desmoplastic stroma encasing tumor nests. Such organization is consistent with a dual biochemical and physical barrier function, with spatial neighbor analyses indicating that fibroblast density peaks at distances coinciding with CD8L T cell accumulation, potentially constraining effector immune cell access to tumor cores.

Despite the generally immune-cold character of HGSOC tumors,^65,66^ spatial profiling revealed recurrent inflammatory hotspots in which malignant cells co-localized with macrophages and T cells. These niches were observed across patients and timepoints, suggesting the presence of persistent but ineffective immune pressure. Similar inflammatory signatures have been reported in single-cell transcriptomic studies across tumor types,^67^ and in ovarian cancer have been coupled to MHC class II–associated programs. Together, these observations support the interpretation that antigen-presenting cells, particularly macrophages, contribute to localized pockets of inflammation. The reproducibility of these niches highlights the value of spatial profiling for resolving immune dynamics that are diluted or obscured by bulk or dissociated approaches.

Consistent with stromal and immune reorganization, CXCL12–CXCR4 signaling was augmented following NACT or PARPi exposure, driven primarily by immune and stromal cells adjacent to tumor clusters. This chemokine axis has been associated with poor prognosis in ovarian cancer,^67,40^ and experimental studies have suggested that its blockade may reduce tumorigenicity.^41,68^ The spatial enrichment of CXCL12–CXCR4 interactions at tumor boundaries, together with treatment-associated differences, supports the view that this signaling axis reflects a property of the remodeled microenvironment rather than a fixed tumor-intrinsic feature.

Neutrophils represented an additional spatially structured component of the post-therapy microenvironment. Tumor-adjacent neutrophils localized preferentially to hypoxic regions and displayed nuclear decondensation patterns consistent with neutrophil extracellular trap (NET) formation. NETs have been increasingly implicated in tumor proliferation, metastasis, and immune suppression.^69–73^ Although functional validation is required, the convergence of spatial localization, morphology, and associated malignant transcriptional states supports the presence of NET-associated niches that may contribute to immune evasion and stress adaptation in treated HGSOC.

Whole-transcriptome profiles complemented the high-resolution insights by enabling inference of large-scale copy number alterations, identification of malignant clones, and prediction of drug sensitivities. Clones were largely patient-specific and conserved across timepoints, consistent with early clonal diversification followed by transcriptional adaptation. Predicted copy number alterations recapitulated known HGSOC driver genes,^3,49^ supporting the validity of the transcriptome-derived inferences. Notably, one clone enriched for HLA-G expression was spatially associated with B cell–rich regions expressing inhibitory receptors, suggesting a localized immune tolerance mechanism. Such mechanisms may represent clinical vulnerabilities, particularly considering that HLA-G-targeted CAR T cells are being evaluated in a phase 3 clinical trial including epithelial ovarian cancer failed or intolerant to platinum therapy and PARPi therapy.^74^

Finally, exploratory drug sensitivity predictions highlighted both expected cytotoxic agents and clinically relevant targeted therapies. A nuclear export inhibitor (KPT-185) emerged among the top candidates, which is approved for multiple myeloma and under evaluation in metastatic ovarian cancer as monotherapy and in combination with paclitaxel and carboplatin. Early-phase trials have reported encouraging activity with manageable toxicity.^75,76^ Additionally, ErbB-associated vulnerabilities were enriched in PARPi-resistant samples, consistent with emerging evidence supporting HER-targeted approaches in select ovarian cancers, including promising responses to trastuzumab deruxtecan in HER2-expressing tumors.^77^ Importantly, spatial variance in predicted sensitivities suggests that such therapeutic response may differ not only between patients, but also across regions or clones within individual lesions, further underscoring how tumor heterogeneity can undermine clinical targeting. Though exploratory, these drug sensitivity analyses illustrate how spatial transcriptomics can prioritize candidate therapies and combinations for functional validation in patient-derived models.

## Conclusions

Overall, our study demonstrates that spatial transcriptomics provide critical insights into how HGSOC tumors and their microenvironments adapt to PARP inhibition. Therapy is associated not only with transcriptional reprogramming, but with coordinated spatial remodeling involving hypoxia, stromal activation, immune exclusion, and localized inflammatory or neutrophil-rich niches. By integrating single-cell resolution with whole-transcriptome coverage, spatial profiling offers a powerful framework for dissecting therapy resistance mechanisms and for generating clinically actionable hypotheses that account for intratumoral heterogeneity.

## Limitations of study

The cohort explored here is small and imbalanced, with limited availability of longitudinal samples from patients who were resistant to initial therapy, complicating interpretation of treatment-specific versus disease-stage effects. Clinical annotation was incomplete for some patients, including HRD status. Technical heterogeneity across samples—including differences in tissue processing (FF vs FFPE), tissue location, and platform resolution—introduces additional variability.

The spot-based resolution of Visium inevitably captures mixed cell populations, particularly in poorly compartmentalized regions, while Xenium’s limited gene panel precludes analyses such as comprehensive pathway inference or CNV estimation and is subject to transcript leakage effects that complicate differential expression testing. Despite these constraints, the concordance of spatial trends across platforms supports the robustness of our core conclusions.

## Supplemental figures

**Figure S2**

A. Top: A bubble plot shows expression of marker genes for each of the granular macrophage populations captured in the Xenium samples. Dot colors correspond to average expression, while dot size corresponds to the percent of cells expressing the gene in each cell state. Bottom: Stacked bars denote the relative contribution of macrophage states relative to all macrophages within each sample.

B. As (A), but for granular T and NK cell populations.

C. The relative abundance change per granular cell type between post- and pre-PARPi samples per patient. Positive values denote enrichment after PARPi, while negative values correspond to higher enrichment in initial timepoint samples. Bars are colored by patient.

D. Heatmap of the top inferred ligand-receptor (LR) activity within RCNs across the Xenium samples. Tiles are colored according to the median percentage of cells with appreciable activity scores across samples for each RCN (x-axis) and LR pair (y-axis).

**Figure S3**

A. Violin plots show the distribution of expression of the indicated gene in malignant cells for each sample, with violins colored by timepoint.

B. A heatmap of the average cancer pathway activity, calculated with progeny and colored according to mean scaled values, within the malignant cells of each sample. Tiles to the left indicate timepoint and patient.

C. Scatter plot of the relationship between patient PFS and the median difference in estimated hypoxia activity in cancer cells between post- and pre-PARPi timepoints of Xenium samples.

D. Heatmap of the pairwise Jaccard similarity of all pseudobulked malignant clusters calculated from the Visium samples. Top annotations denote cluster clade groupings, sample timepoint, and patient. Tiles are colored by Jaccard similarity of malignant clusters’ marker genes. Boxes below show the top consistent genes for clusters in clades 3, 4 and 5.

**Figure S4**

Line plots highlight the relationship between minimum distance to a cancer cell and the total number of neighbors in fibroblasts (left), the number of fibroblast neighbors in fibroblasts (middle) and the number of fibroblast neighbors in CD8+ T cells (right) within a 25µm radius. Lines are colored by patient (pre-PARPi only) and the gray regions show confidence intervals.

**Figure S5**

A. A bubble plot shows the CXCL12-CXCR4 activity within TME cell types of the Xenium samples. Bubbles are colored by the average activity scores (non-zero cells only) and sized according to the proportion of cells with appreciable values.

B. Violin plots show the expression of neutrophil extracellular trap genes included in the Xenium panel (*MPO*, *ELANE*, *PADI4*) per sample, with distributions colored by timepoint where applicable.

C. For the four Xenium samples in which neutrophils were detected, stacked bars show the relative contribution segmentation method used to define cell boundaries by Xenium across cell types. Neutrophils and LQ cells are highlighted.

D. Neutrophil co-localization with other immune and stromal cell types across samples for which neutrophils were detected. Tiles are colored by scaled likelihood of cell types being within 50 µm.

**Figure S6**

A. Density plot of the reference-based ovarian cancer cell proportion per spot for Vis-9-CT-A. The dashed line corresponds to the OTSU cutoff, and the background is colored according to how spots in those regions were classified.

B. Boxplots of the reference-free proportion estimates for stroma, immune and ovarian (OV) epithelial cells colored by the reference-based classification labels assigned for all Visium samples.

**Figure S7**

A. Scatterplot of the relationship between the total number of malignant spots (x-axis) versus the number of inferred CNV clones (y-axis) for each Visium sample. Points are colored by patient and shaped by treatment timepoint.

B. Bar plots indicate the most recurrent cancer driver genes predicted to be amplified (top) and deleted (bottom) in Visium samples, with the x-axis denoting the number of samples in which at least one clone harbored the event and the y-axis indicating the gene.

C. Heatmaps show the full predicted CNV profiles of clones from patients with top (left) and longitudinal (bottom) samples.

## Methods

### Human tissue samples

Human tissue samples were obtained from ovarian cancer patients following informed consent in accordance with ethical and legal guidelines outlined in the Declaration of Helsinki. The study protocol iPARPresist used was reviewed and approved by the Ethical Committee of the University Hospital Germans Trias i Pujol (HUGTP), under the approval number PI19-190.

### Sample tissue preparation

#### Visium v1 whole-transcriptome Fresh-frozen samples

HGSOC samples were embedded in cryomolds using OCT (Tissue Tek Sakura) using a bath of isopentane and liquid nitrogen (Tissue preparation guide CG000240, 10x Genomics). Following freezing in OCT, blocks were stored at -80°C. RNA integrity was assessed by calculating RNA Integrity Number (RIN) of freshly collected tissue sections with Qiagen protocol (RNeasy Mini Kit 74104) and analyzed by TapeStation. Samples with Rin above 7 were selected for the experiments. For sectioning, blocks were equilibrated to -20°C in the cryochamber (LEICA CM1950). 10 µm-thick sections were placed onto the active areas (6,5 mm x 6,5 mm) of chilled 10x Genomics Visium Slides.

For Tissue Optimization, 7 sections were placed on a Tissue Optimization Slide (3000394, 10X Genomics) then fixed in chilled methanol and stained according to the Visium Spatial Tissue Optimization User Guide (CG000238, 10X Genomics) to determine optimal permeabilization time for HGSOC Tissue. For tissue optimization experiments, fluorescent images were taken with a TRITC filter using a 10X objective and 900 ms exposure time. Optimal Permeabilization time was determined at 30min of incubation.

For Gene Expression experiments, 4- 10 µm-thick sections were placed onto Gene Expression slide (2000233, 10X Genomics) and adhered by warming the back of the slide with the tip of the finger. Gene Expression slide was kept at -80 °C for 1 week.

On the day of the experiment, tissue sections were fixed in chilled methanol at -20 °C for 30 minutes, and Hematoxilin & Eosin staining was performed according to the Demonstrated Protocol (CG000160, 10x Genomics). Brightfield histology images were taken using a 10X objective (Plan APO) on a Nikon Eclipse Ti2, images were stitched together using NIS-Elements software (Nikon) and exported as tiff files. Following imaging, the coverslip was removed gently, and samples were permeabilized for 30 minutes. cDNA Synthesis and amplification was performed following Visium Spatial Gene Expression User Guide (CG000239, 10X Genomics).

Libraries were prepared according to the Visium Spatial Gene Expression User Guide (CG000239, 10X Genomics) and sent for sequencing. Sequencing depth was calculated with the formula (Coverage Area x total spots on the Capture Area) x 50,000 read pairs/spot.

Sequencing was performed using the following read protocol: read 1: 28 cycles; i7 index read: 10 cycles; i5 index read: 10 cycles; read 2: 90 cycles.

#### Visium v1 whole-transcriptome Formalin-Fixed, Paraffin-Embedded samples

FFPE samples were placed in the microtome and sectioned 7 µm thick. After floating on a water bath at 42 °C, sections were placed on Visium Spatial Gene Expression slides (2000233, 10X Genomics). Afterwards, the slides were dried at 42°C for 3 hours. The slides were then placed inside a slide mailer, sealed with parafilm, and left overnight at Room temperature.

Deparaffinization was performed by successive immersions in xylene and ethanol followed by H&E staining according to Demonstrated Protocol (CG000409, 10X Genomics).

Brightfield images were taken using a 10X objective (Plan APO) on a Nikon Eclipse Ti2, images were stitched together using NIS-Elements software (Nikon) and exported as tiff files. After imaging, the glycerol and cover glass were carefully removed from the Visium slides by holding the slides in an 800 ml water beaker and letting the glycerol diffuse until the cover glass detached and density changes were no longer visible in the water. The slides were then dried at 37°C.

Libraries were prepared according to the Visium Spatial Gene Expression for FFPE User Guide (CG000407, 10X Genomics) and sent for sequencing Using HiseqX 150PE (2x 150bp) applying 1% Phix. Sequencing was performed using the specific protocol for Visium v1 FFPE read protocol: read 1: 28 cycles; i7 index read: 10 cycles; i5 index read: 10 cycles; read 2: 50 cycles.

#### Xenium Prime 5K sample preparation

The Xenium Prime workflow began by sectioning 5μm FFPE tissue sections onto a Xenium slide, according to the "Xenium FFPE Tissue Preparation Guide" (CG000578 Rev C, 10X Genomics). Briefly, the sections were cut, floated in an RNAse-free water bath at 42 LJC, and carefully placed onto the capture area of a Xenium slide (PN- 3000941). After sectioning, the slides were incubated at 42 LJC per 3 hours and kept in a sealed bag with desiccators at room temperature overnight.

The next day, the slides were incubated at 60 degrees for 30 minutes and the slides were processed following the "Xenium In Situ for FFPE- Deparaffinization and Decrosslinking" protocol (CG000580 Rev D, 10X Genomics). After deparaffinization, the Xenium slides were assembled into Xenium cassettes, which allow for the incubation of slides on the Xenium Thermocycler Adapter inside a Thermocycler machine with a closed lid for optimal temperature control, and Decrosslinking step was performed.

The slides were then processed using the “Xenium Prime In Situ Gene Expression protocol” (CG000760 Rev A, 10X Genomics) with Cell Segmentation reagents.

Briefly, after Priming hybridization and Rnase treatment, the samples were incubated at 50 LJC overnight for approximately 19 hours with the Xenium Prime Human Panel which targets 5000 human genes. This was followed by a series of washing steps, a ligation step, and amplification. Afterward, the slides were treated with an autofluorescence quencher and a nuclei staining.

Finally, at the end of the third day of the protocol, the slides were loaded into the Xenium Analyzer. The first step in the Xenium instrument consists in sample scanning, where images of the fluorescent nuclei in each section are given, and these images allow the user to select and determine the regions to be included in the analysis.

After selecting all samples, the run in the Xenium Analyzer lasted 7 days. The instrument was emptied of consumables, and the Xenium slides carefully removed. PBS-T was added to the slides and the slides were stored at 4 LJC for 1 week.

### Post-Xenium Analyzer H&E staining

Slides were processed following post-xenium analyzer H&E protocol (CG000613 Rev A, 10X Genomics). A quencher removal solution was prepared immediately before use by dissolving sodium hydrosulfite in Milli-Q water to a final concentration of 10 mM. The slides were gently immersed in the quencher removal solution using a slide mailer and incubated for 10 minutes at room temperature. This was followed by three washes in Milli-Q water.

The sections were then stained with Gill’s hematoxylin for 20 minutes, followed by three washes in Milli-Q water. Bluing solution was applied for 1 minute, followed by a Milli-Q water wash. The slides were subsequently dehydrated through graded ethanol solutions (70% and 95%), counterstained with alcoholic eosin for 2 minutes, further dehydrated to xylene, and finally mounted with coverslips.

Brightfield histology images were taken using 10X and 20X objectives (Plan APO) on a Nikon Eclipse Ti2, images were stitched together using NIS-Elements software (Nikon) and exported as tiff files.

### Xenium computational analyses

#### QC and pre-processing

Gene expression profiles matrices from Xenium Analyzer (v3.0.0.15) were loaded with Seurat and filtered to retain only cells with >= 10 transcripts. Each sample’s data was normalized and scaled using `SCTransform(return.only.var.genes = FALSE, conserve.memory=TRUE, vst.flavor = "v2")`. A principal component analysis (PCA) was conducted on the top 1500 highly variable genes (HVGs), and PCs 1:30 were used to run UMAP. Unless otherwise stated, the SCT assay was used in downstream analyses involving gene counts, with the exception of the sketch approach for cell typing and the determination of cell-specific markers, both which relied on log-normalized counts.

#### Cell type annotation and longitudinal differences

To conduct initial cell typing per sample, we leveraged Seurat’s sketch approach. After filtering, the data was log normalized and scaled before determining the top 2000 HVGs. The 50K cells contributing to the variable gene signal were selected by LeverageScore and subset. On this subset, it was run PCA, UMAP, and Louvain clustering at a resolution of 1.0. The remaining cells were projected onto the sketched data using the sketched UMAP representation. The marker genes were determined for each cluster using `FindAllMarkers()`. The cell type gene lists from CellMarker 2.0^78^ were utilized to run over representation analysis for each cluster’s marker gene list, defining the 5000 Xenium gene panel as the universe with clusterProfiler’s `enricher(pvalueCutoff = 0.05, qvalueCutoff = 0.05)`. In tandem, UniCell Deconvolve^79^ was also run on each sample’s data, and the top cell annotations for each cluster were defined by the highest mean cell type score.

Using cluster marker genes, enriched cell types from CellMarker 2.0, and the UCD top cell annotations for cells in each cluster, we manually annotated initial cell types with general labels. A subset of cells displayed ambiguous phenotypes due to low transcript counts or mixed signals. These were retained in spatial analyses as low-quality cells (LQ), as they were consistently distributed throughout the tissue and unlikely to be artifacts. At this stage, we also both used both the hepatocyte signature and pathologist annotations to exclude the liver area/hepatocyte cluster cells from sample Xen-3 postPARPi. Once initial general cell types were assigned, the cell-type specific markers for each sample were determined with `FindAllMarkers(only.pos = T, min.pct = 0.1)`.

Refinement of each of macrophages, T cells, and B cells, and vascular cells (those annotated as “endothelial” or “pericyte”) were done by subsetting cells with these labels in each sample, rerunning sketch, and integrating across samples. In each cell-specific subsets, we selected up to 5000 cells from each sample via the LeverageScore. The top 2000 HVGs were determined, and to limit the influence of potential transcript leakage on downstream clustering, we excluded previously determined specific markers of other cell types from the HVG list (genes with average log2 fold change > 2, with the exception of DC markers in the evaluation of macrophages due to functional similarity).The refined HVG set was then used to run PCA and integrate cells between samples with CCA.

Each integrated dataset was subject to Louvain clustering at resolutions of 0.2, 0.5, 1.0 and 2.0 before determining marker genes for each. The cluster marker genes were used to select a resolution for subtype separation and for annotation of granular populations (ex. only resolution 2.0 separated out *RORC* expressing Th17 cells for the integrated T cells). The cell subtypes were then assigned to the appropriate cluster before sketch-derived clusters were projected onto the full dataset and the granular cell labels were mapped back to samples.

Differences in cell type proportion abundance for each granular cell type between prePARPi and postPARPi samples were computed as the logit-transformed differences within each patient (Fig S2c).

#### Defining recurrent cellular neighborhoods

Recurrent cellular neighborhoods (RCNs) were identified using the Scimap package. For each cell, we quantified the composition of neighboring granular cell types within a 50 µm radius by calculating the fraction of each cell type surrounding a given center cell. These neighborhood profiles were then grouped using k-means clustering (k = 20) across all six Xenium samples using the function `sm.tl.spatial_cluster(r=50, k=20).` The resulting RCNs were annotated manually based on the relative abundance and enrichment patterns of cell types within each cluster.

#### Ligand/receptor activity estimation

The list of Ligand-Receptors (LR) used to score in the Xenium dataset was obtained from CellChat^80^. The stLearn package was used to score the LR pairs by identifying cells expressing the ligand and neighboring cells expressing the corresponding receptor within a 30 µm radius. SCT-normalized gene expression values were used as input. Statistical significance of the LR interaction scores was assessed using 10,000 randomly generated cell pairs. `st.tl.cci.run(min_spots = 20, distance=30, n_pairs=10000)`

To handle large tissue areas, the tissue was subdivided into spatial batches of 2000 µm × 2000 µm, with an additional 30 µm buffer region surrounding each batch. The buffer region was included during score computation to avoid edge effects, but scores originating within the buffer were discarded. Finally, scores from all batches were aggregated.

#### Ligand/receptor activity within recurrent cellular neighborhoods

To evaluate whether certain LR might be enriched in particular RCN, we ran per-sample Wilcoxon signed-rank tests to evaluate the scores of each LR pair in a particular RCN versus all other RCNs. In addition to the test p-value, we also noted the mean LR scores in each group (RCN X versus all others), the median in non-zero cells, and the proportion of cells in each group with non-zero scores. P-values for results per sample were adjusted with FDR.

Due to the large amount of dropout in LR values across cells, the proportion of cells with a non-zero value in the RCN test groups were leveraged for filtering. To select the top LR pairs for plotting, the results per sample were filtered by those with Wilcoxon FDR less than 0.001 and a difference larger than 0.1 between the proportion of cells in each group. The top 10 LR per RCN in each sample, defined by group proportion difference, were selected. For final plotting only top LR per RCN repeated in multiple samples and with a median percentage of cells expressing across samples larger than 5% were kept (Fig. S2d).

#### Bulking and pseudobulking transcripts

To overcome the issue of sparsity posed by Xenium data, we subjected the sample counts to bulking and pseudobulking procedures. To generate sample-level pseudobulk data, raw gene counts from cells of interest (e.g., fibroblasts) were aggregated per sample. Variance stabilizing transformation (VST) was applied to the resulting pseudobulk counts for dimensionality reduction using PCA. Differential expression analysis between timepoints was conducted on pseudobulk counts using “patient” as a cofactor. Log2 fold changes were estimated and shrinkage was applied using the *apeglm* method via `lfcShrink`.

Pseudobulking herein refers to rasterizing the single-cell data into "pixels". To do this, the sf package was used to generate a grid of 55um diameter hexagons overlaying the tissue, `st_make_grid(cellsize = 55, square = FALSE)`. In the event that a cell fell on the border of the grid and was assigned to multiple pixels, only its first appearance was retained. The counts of the cells assigned to each of the pixels were aggregated and log normalized with Seurat’s AggregateExpression function. For each generated pixel, the contributions of each cell type were retained as meta data. In addition to cell-agnostic aggregation, this approach was orthogonally done in a cancer-focused way (“cancer raster”), in which pixels with at least 3 cancer cells and 70% contribution by cancer cells were subset and counts just from the malignant cells in each pixel were aggregated.

#### Cancer sub-clustering

Using the cancer raster data, we generated expression-based clusters of the malignant areas in each sample. The previously log-normalized data was scaled and subject to PCA and Louvain clustering, during which genes that were previously determined as markers for other cell types were excluded from the HVG set. Marker genes were determined for each of the subclusters in each sample with `FindAllMarkers(only.pos=TRUE, min.pct= 0.1)`.

Cancer raster cluster marker genes with adjusted p-value less than 0.001 and an average log2 fold change greater than 0.5 were retained as positive markers and used to compute the Jaccard similarity index for all pairwise cancer raster clusters in all samples, defined as:

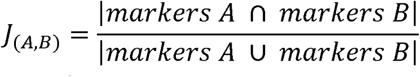

Where, if A and B is a given pair of clusters, J_(A,B)_ is the Jaccard similarity between clusters A and B, calculated by dividing the number of intersecting marker genes between clusters A and B by the union of marker genes from clusters A and B.

The Jaccard similarity matrix between all pairwise cluster combinations was then clustered using euclidean distance and `hclust()` to generate 5 clades of similar clusters across samples. Top recurrent marker genes for the clusters in each clade were determined.

#### Protein intensity scoring

To quantify Vimentin and αSMA signal intensity at the single-cell level, channel *ch0003* from the Xenium multimodal segmentation output and the associated cell segmentation boundaries were imported into QuPath^81^. Mean pixel intensity was calculated for each individual cell by averaging pixel values within the corresponding cell boundary.

#### Pathway scoring

Gene set enrichment analysis was performed using fgsea on bulked gene expression profiles from cancer cells in each sample. Counts were aggregated and normalized using DESeq2 size factor normalization without differential expression modeling. Log2 fold changes between conditions were calculated per patient from normalized counts with a pseudocount of 1, and genes were ranked by the product of log2 fold change and log-transformed mean expression. Enrichment analysis was performed using MSigDB Hallmark gene sets, with results computed independently per patient.

The transcriptional footprint of Hypoxia and other canonical signaling pathway activities was extracted with PROGENy^29^ using the top 100 genes for each pathway. The pathway activity was inferred using a multivariate linear model implemented in decoupleR^82^ using the SCT-normalized gene expression at the single-cell level.

In addition to scoring curated pathways, we also used a data-driven approach to define the an interferon (IFN) response signature consistently observed in the Xenium samples. In the initial sample sketch clusters, multiple samples exhibited clusters defined by genes related to IFN induction, such as of *IFIT* genes 1-3 and *CXCL*s 9-11. We defined a comprehensive signature by compiling the marker genes for these clusters (Bonferroni adjusted p-value < 0.01 and average log2 fold change > 1) and further requiring that the gene be repeated in at least four out of the 5 total clusters with IFN response signatures, resulting in 37 genes (Table S6). Individual cells and rasterized pixels in each sample were scored with UCell’s AddModuleScore_UCell.

#### Calculating pairwise distance between cells

For every cell in each sample, the minimum distance to each cell type present (i.e. Cancer) was determined using `KDTree` (scipy.spatial). Each cell’s x and y coordinates and granular cell type label was used to query the nearest neighbor of a given cell type within a maximum search radius of 500 µm. Self-matches were removed and, in the event that no cell of a given type was identified within the radius, the distance was set to 501 µm and the neighbor’s ID was set to a missing value.

#### Modeling gene expression changes with decay functions

Distance-dependent gene expression was modeled with generalized additive models (GAMs) with thin plate regression splines, using gene expression from a given cell type (e.g., cancer cells) as a function of the minimum distance to another cell type (e.g., fibroblasts). Models were fitted using the bam function (mgcv) with a Gaussian family. Predicted expression values were generated across the observed distance range, with peak expression defined near zero distance and basal expression estimated from distal regions. Expression profiles were normalized between peak and basal levels, and decay metrics D50 and D90 were computed as the distances at which expression decreased to 50% and 10% of the peak amplitude, respectively. Genes with a short D90 distance (< 50 µm) were considered to exhibit leaked gene expression from neighboring cells.

#### Quantifying T cell infiltration

To numerically define the extent of T cell infiltration into malignant areas, we leveraged the RCN assignments for each cell. For each sample, we quantified the total number of T cells present in the sample outside of TLS regions (to limit bias that would result in an inflated denominator) by summing all T cells but those in RCNs 18-20. We then defined malignant-infiltrating T cells as those present in cancer-dominant regions by summing all present in RCNs 1-3. For each sample, we divided the number of malignant infiltrating T cells by the total number of non-TLS T cells and compared the values per patient between timepoints (Fig. 5a).

#### Cell co-localization in raster pixels with high IFN response signature

To determine whether certain cell types tended to co-localize more frequently in sample areas with high IFN signature scores, the rasterized data was leveraged. First, pixels were annotated as IFN high if they scored the top 10% of all IFN scores in that sample. For each possible pair of cell types in a given sample, we created a 2x2 contingency table of the co-occurrence of the two cell types against IFN status and applied Fisher’s exact test to estimate an odds ratio (OR) and p-value describing whether that cell-type pair co-occurs more frequently in high-IFN pixels than in low-IFN pixels. The log2 OR was then computed as an effect size measure.

To prioritize consistent cell pairs co-localized across samples, we computed the mean log2 OR across samples and counted the number of samples in which each pair was observed. Cell-type pairs detected in more than two samples and with a mean log2 OR greater than 1 were retained for plotting in an undirected network graph with `ggraph`, visualized using a force-directed (Fruchterman–Reingold) layout (Fig 5d).

### Visium computational analyses

#### QC and pre-processing

For Visium data generated in this study, FASTQ files were processed with SpaceRanger v2.0.0 and the GRCh38-2020-A reference. Sample count data was read with Seurat, and for the FF samples, genes were filtered to include only those in the FFPE probe set (v1.0) before re-calculating the number of genes and counts for each spot. Spots with less than 750 genes were filtered out, and any remaining spots not on the bulk tissue region were manually excluded. Retained spots in each sample were normalized and scaled using `SCTransform(vst.flavor= "v2", method = "glmGamPoi", return.only.var.genes = F)` before running PCA.

A previously published HGSOC single-cell dataset, the MSK SPECTRUM cohort^9^, was leveraged for reference-based deconvolution. The processed full cohort data was downloaded from synapse and the raw counts and metadata therein were used to make a clean Seurat object to ensure version compatibility. Long non-coding RNA were filtered out from the features, as well as cells from ascites samples and those ascribed to “Other” in the cell type assignments. The full dataset was downsampled in a patient-balanced manner to retain a maximum of 10 thousand cells per cell type. We did this 1) to lessen computational load and 2) to minimize the odds that a given cell type would be biased by contribution from patients with a substantial number of those cells present. Namely, instead of blindly selecting 10,000 random cells per cell type, we restricted the number of cells that could be selected per patient to ensure capture of gene expression profiles across individuals. The resulting downsampled data was not normalized, as it was used to run a deconvolution method that leverages raw counts.

Finally, we also downloaded and processed a healthy ovarian Visium dataset^48^ for use as a healthy reference when estimating CNV events. We first excluded Visium samples from young (18–28 years) individuals, as these were dissimilar to our patients’ age profiles (Table S1). Then, we created Seurat objects from each remaining sample’s count data, filtering the genes to include only those in the Visium probe set retained in our samples, and similarly excluding spots with less than 750 genes. Spots in each sample were normalized and scaled using `SCTransform(vst.flavor= "v2", method = "glmGamPoi", return.only.var.genes = F)` before running PCA and using the first 30 components to determine neighbors and run Louvain clustering (default 0.8 resolution). The resulting cluster assignments were retained for each sample for downstream CNV analysis.

#### Spot deconvolution and classification

First, we mapped the expression profiles from the filtered, downsampled single-cells published in the MSK SPECTRUM cohort^9^ to each Visium sample with Robust Cell Type Decomposition^83^ (RCTD) using the “full” mode with `run.RCTD()`. The resulting weights per spot were normalized with `normalize_weights()`. Within each sample, we then applied Otsu’s method {https://doi.org/10.1109/TSMC.1979.4310076} to the distribution of normalized ovarian cancer cell estimates across all spots to binarize the spots into two categories. Then, we added a third “buffer” category to account for mixed spots according to a margin of 0.05 above and below the threshold value, resulting in three classes: TME, buffer (mixed), and malignant (Fig. 6b and S6a).

We also used a reference-free deconvolution tool, UniCell Deconvolve (UCD)^79^, a deep learning model pre-trained on over 28 million cells across hundreds of cell types. The SCTransformed data per sample was output to a matrix, alongside cell names and coordinates, to make AnnData objects in python. The tool was applied to each sample with <u>ucd.tl</u>.base(verbosity=logging.DEBUG, split=True, propagate=True, use_raw=False) and the resulting predictions for each spot were saved. Using the cell types detected in the Xenium samples as a guide, we selected the most relevant cell types from UCD (Table S7).

After subsetting the estimates for these cells, we zeroed out any contribution below 0.05 and scaled the estimates to sum to 1 for each spot. We then summed the UCD estimates of stromal and immune cell types (Table S7) and retained the ovarian epithelial estimates for each spot for comparison with the RTCD-based classifications used in downstream analyses (Fig S6b).

#### Malignant entropy and dispersion estimates

We quantified spatial dispersion of malignant spots using Marcon and Puech’s *M*^84^ via the dbmss R package^85^. For each malignant spot, *M* measures the local proportion of malignant neighbors within a given radius and normalizes this quantity by the global malignant spot proportion in the sample.This yields *M(r)*, where *M(r)* =1 indicates spatial randomness, *M(r)* >1 indicates local clustering, and *M(r)* <1 indicates spatial avoidance.

For each sample, we generated *M*-envelopes in 100 µm increments up to 2000 µm under a random location null model with MEnvelope(r=seq(0,2000,100), NumberOfSimulations=100, Alpha=0.05, ReferenceType="malignant", CaseControl=FALSE, SimulationType="RandomLocation", Global=FALSE). We then extracted *M*(r= 400μm) per sample as a summary measure of malignant self-dispersion. For simpler visualization, the *M* values were transformed with log10 before plotting (Fig. 6c).

We also calculated cell-type mixing in malignant spots by calculating the Gini-simpson entropy. For each spot classified as “malignant”, we pulled the RCTD-derived cell type proportion estimates. Entropy was estimated per-spot as the following:

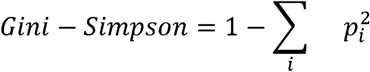

Wherein *p_i_* denotes the deconvolution-estimated proportion of cell type *i*, where i ∊ {i, …,k} Higher values indicate greater cell-type diversity within a spot, while lower values represent a purer malignant signature. Resulting entropy values were averaged per sample for comparison (Fig. 6X).

#### Distances between malignant and immune regions

We determined the nearest distance between each malignant spot and an immune-enriched spot in each sample. First, aggregate immune contributions per spot were determined by summing RCTD estimates of all immune cell types (i.e., all cell types except endothelial cells, fibroblasts, and ovarian cancer cells) to obtain a composite score. Spots that were not previously annotated as malignant and that had summed immune contribution greater than 0.25 were retained for distance measures.

We then used the spot coordinates (in µm) to compute the pairwise Euclidean distances between all malignant and immune-enriched spots with the `rdist` function from the `fields` R package. For each malignant spot, we identified the nearest immune-enriched spot by minimum distance and recorded both the distance and the barcode of this nearest neighbor.

#### Malignant CNV inference

For each sample, we subset the spots assigned to the malignant compartment to estimate CNV profiles from the expressions data with InferCNV^86^. We then merged the subset object with the healthy ovarian reference Visium samples^48^ to overcome limitation of few healthy spots/extensive mixing in some samples that might bias CNV calling. To select the best areas of the healthy reference to define a background signature, we used a data-driven approach. We aggregated expression according to sample clusters, combining expression in all malignant spots from our sample and within previously-determined Louvain cluster assignments in healthy samples. After determining HVGs, running PCA, and calculating the nearest neighbors, we subset the 10 nearest healthy cluster neighbors to use as reference in InferCNV.

Objects for running InferCNV were made using the counts from malignant spots and those belonging to the neighboring reference clusters previously determined. For gene ordering, GENCODE Release 27 (GRCh38.p10) was used. The mitochondrial chromosome was excluded, but the X chromosome was retained given that all samples were of biological female subjects.

Per sample, we ran InferCNV with leiden resolution 0.001 up to step 15 and assessed resulting subclusters: infercnv::run(cutoff=0.1, cluster_by_groups=F, denoise=T, plot_steps=T, HMM=T, analysis_mode = "subclusters", tumor_subcluster_partition_method="leiden", leiden_resolution= 0.001, up_to_step =15, scale_data=T). In instances that seemed to lack groupings of spots with unique events, we reran the algorithm increasing resolution to 0.005, 0.01, 0.05 and 0.1 where necessary. If groups did not separate well at the subsequent resolution, the previous was kept. Samples with resolution deviations were: Vis-8-PARPi with 0.05, Vis-10-Dx with 0.01, Vis-11A-CT with 0.05, Vis-11B-CT with 0.01 (groupings did not improve for Vis-10-Dx nor Vis-11B-CT at 0.05 resolution; similar for Vis-8-PARPi and Vis-11A-CT at 0.1 resolution).

After determining the optimal resolution for each sample, the full pipeline was run by omitting the `up_to_step` argument. To get the gene-level predictions for each sample, the HMM_CNV_predictions.HMMi6.leiden.hmm_mode-subclusters.Pnorm_0.5.pred_cnv_genes.dat file was used. Genes with an inferred state = 3 were classified as neutral, while those with state > 3 were predicted to be amplified and those with state < 3 were predicted to have undergone deletion. In resulting clone assignments of spots, we filtered out any clones with only 1 spot assigned and denoted these as “singletons” in plots. In subsequent clone-based analyses, these singletons were omitted.

#### Clone CNV similarity

To determine similarity between inferred CNV clones, we calculated the Jaccard similarity between CNV profiles. Per clone, we filtered out neutral status genes and constructed a clone-by-gene matrix where each entry represents the CNV status of a given gene in that clone. CNV states were collapsed into a ternary encoding in which amplifications (state > 3) were set to 1, deletions (state < 3) were set to –1, and genes without a CNV event remained 0 (if present in another clone as a predicted CNV event). This yielded a discrete CNV signature vector for every clone across the union of all genes.

We then computed a Jaccard similarity index for every pair of clones based on their non-neutral CNV profiles. For a given pair of clones, the intersection was defined as the number of genes for which both clones shared the same non-zero CNV state (i.e., both amplified or both deleted). The union was defined as the total number of genes with a non-zero CNV state in at least one of the two clones. The Jaccard similarity for each given pair of clones A and B was calculated as:

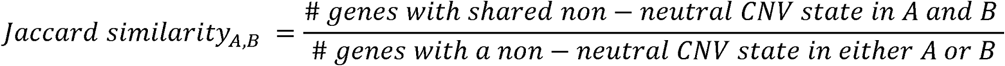

Clone identifiers were parsed to recover the originating sample for each clone, and these were merged with clinical metadata to annotate the patient and clinical timepoint associated with each clone. Each clone pair was then classified into one of four relationship categories: same (clones from the same sample), consecutive (clones from different samples of the same patient collected at the same clinical timepoint), longitudinal (clones from different samples of the same patient collected at different timepoints), or unrelated (clones from different patients).

The resulting table of pairwise clone–clone Jaccard similarities, together with their sample, patient, and relationship annotations, was exported for downstream analysis. Distributions of Jaccard similarity across relationship categories were visualized using violin plots (Fig. 7a).

#### Cancer driver genes affected by CNVs

We filtered the CNV-affected genes by known cancer driver genes using a previously published gene list^87^. We separately assessed the most common drivers predicted to be amplified and deleted by counting the event once per sample occurrence (rather than once by clone to prevent bias from samples with more clones). The top drivers predicted to be altered by amplification or deletion across the Visium samples were selected for plotting.

#### Clonal differential expression analyses

To determine how gene expression patterns might associate with clonal assignments, we annotated spots according to their clone assignments (excluding singletons) and “Buffer/TME” for non-malignant spots. We then used `FindAllMarkers` from Seurat to determine differentially expressed genes between groups. Results for each sample are detailed in Table S5.

#### Drug sensitivity estimations

Spot-level drug sensitivity estimates were performed using BeyondCell^53^, which uses predefined pharmacogenomic gene sets to compute drug response signatures. Drug response signatures were obtained from the SSc (sensitivity score) collection included in BeyondCell, which contains curated gene sets associated with compound sensitivity profiles, using `GetCollection(SSc, include.pathways = TRUE)`.

For each sample, BeyondCell Scores (BCS) were calculated using `bcScore` with a minimum gene expression threshold of 0.1. The BS represent normalized enrichment estimates of drug sensitivity at the spot level across the SSc gene sets. Drug prioritization analyses were then performed using `bcRanks`, grouping spots by their RCTD-derived class labels to rank compounds by their relative sensitivity across groups by aggregating BCS values, enabling identification of drugs selectively predicted to be effective in malignant regions versus non-malignant (buffer/TME) compartments.

The resulting drug ranking table and the full BeyondCell object (including normalized and scaled BCS matrices) for each sample were exported for downstream analyses and visualization. Drug set metadata provided by BeyondCell were also retained for interpretability.

