## Supplementary figures and images for "Convergent stromal and immune remodeling defines spatial tumor dynamics in PARP inhibitor–resistant high-grade serous ovarian cancer"

### Supplemental figures

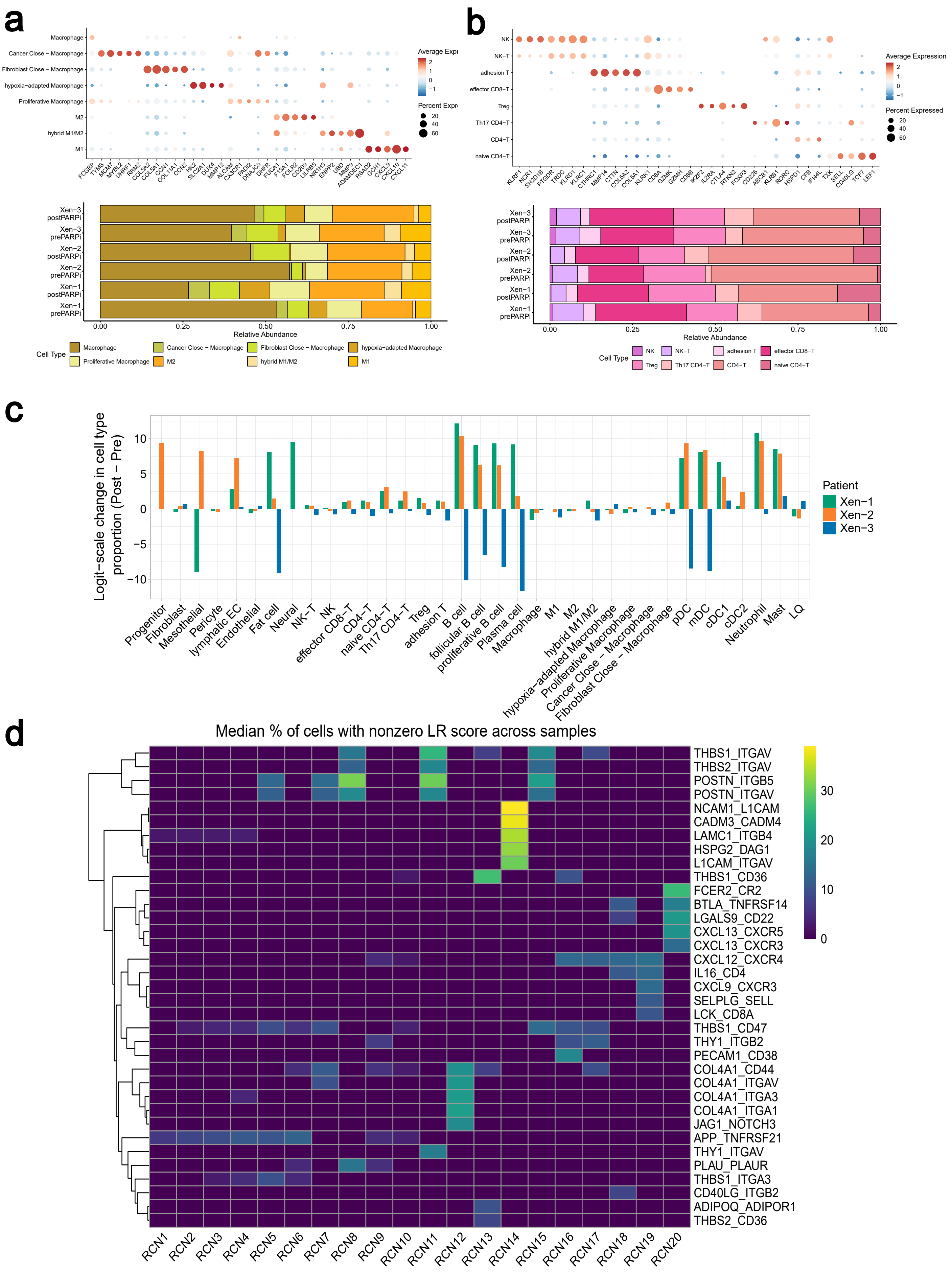

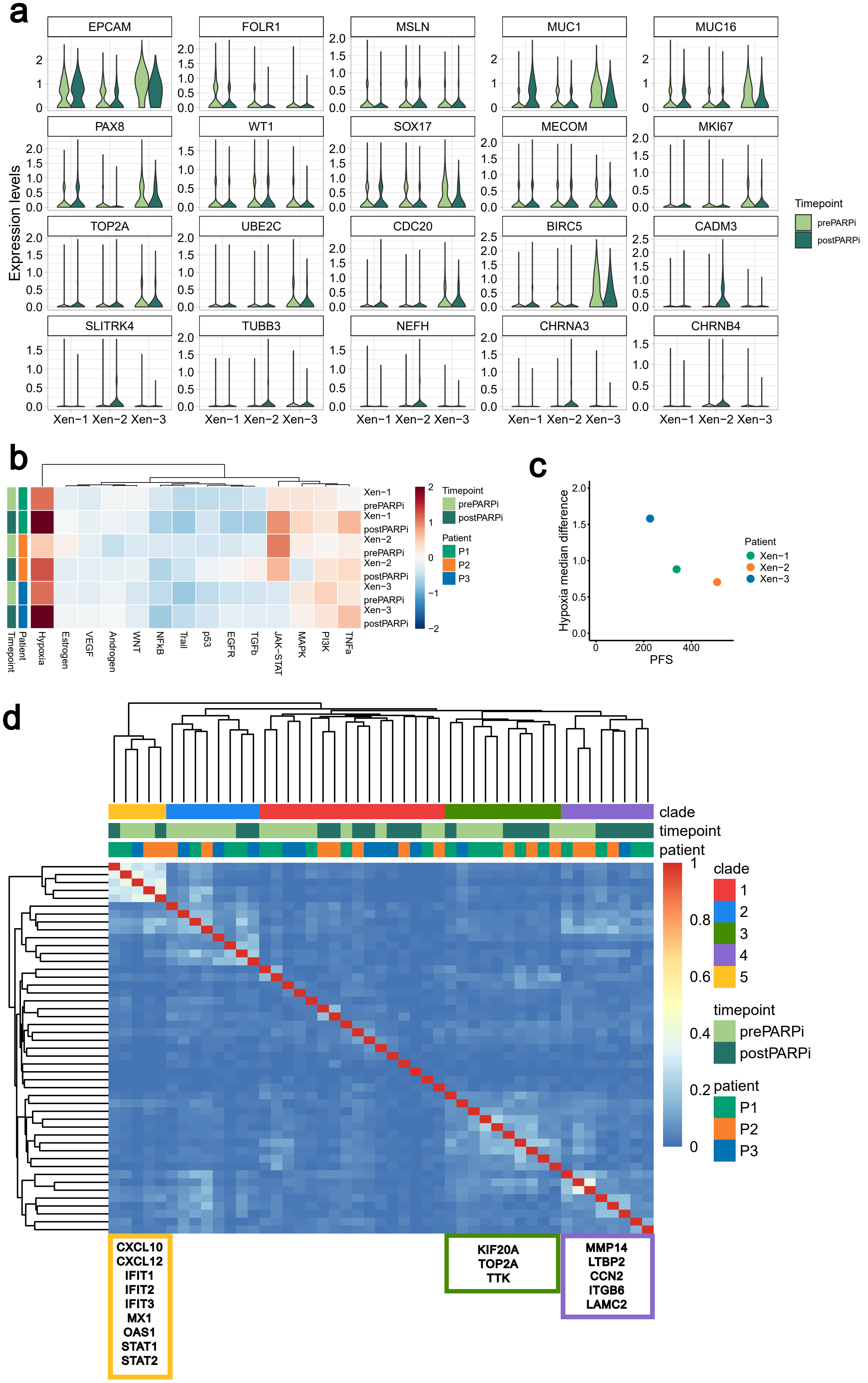

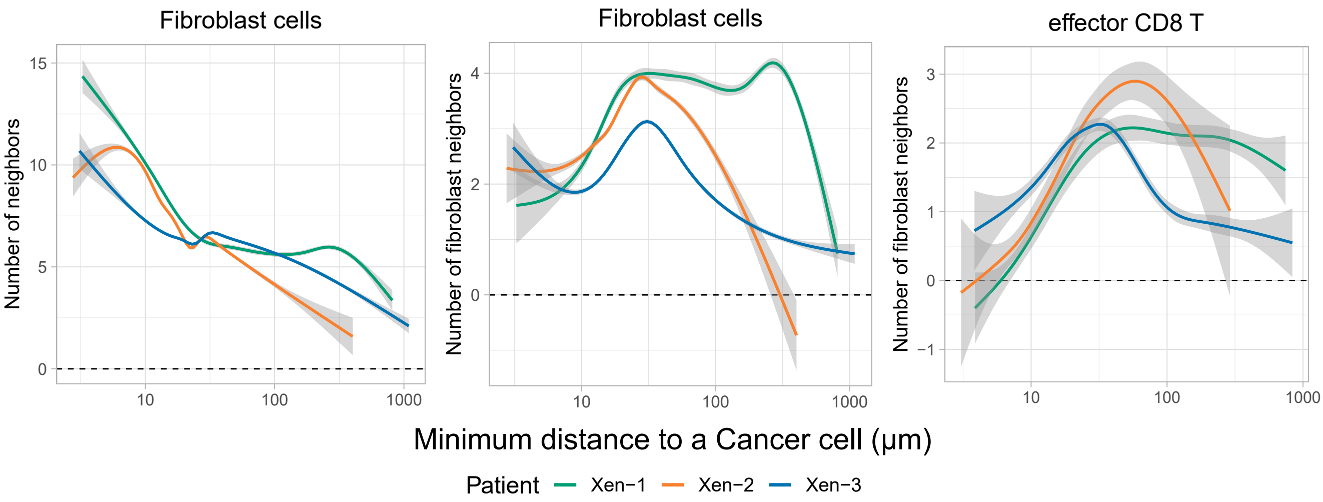

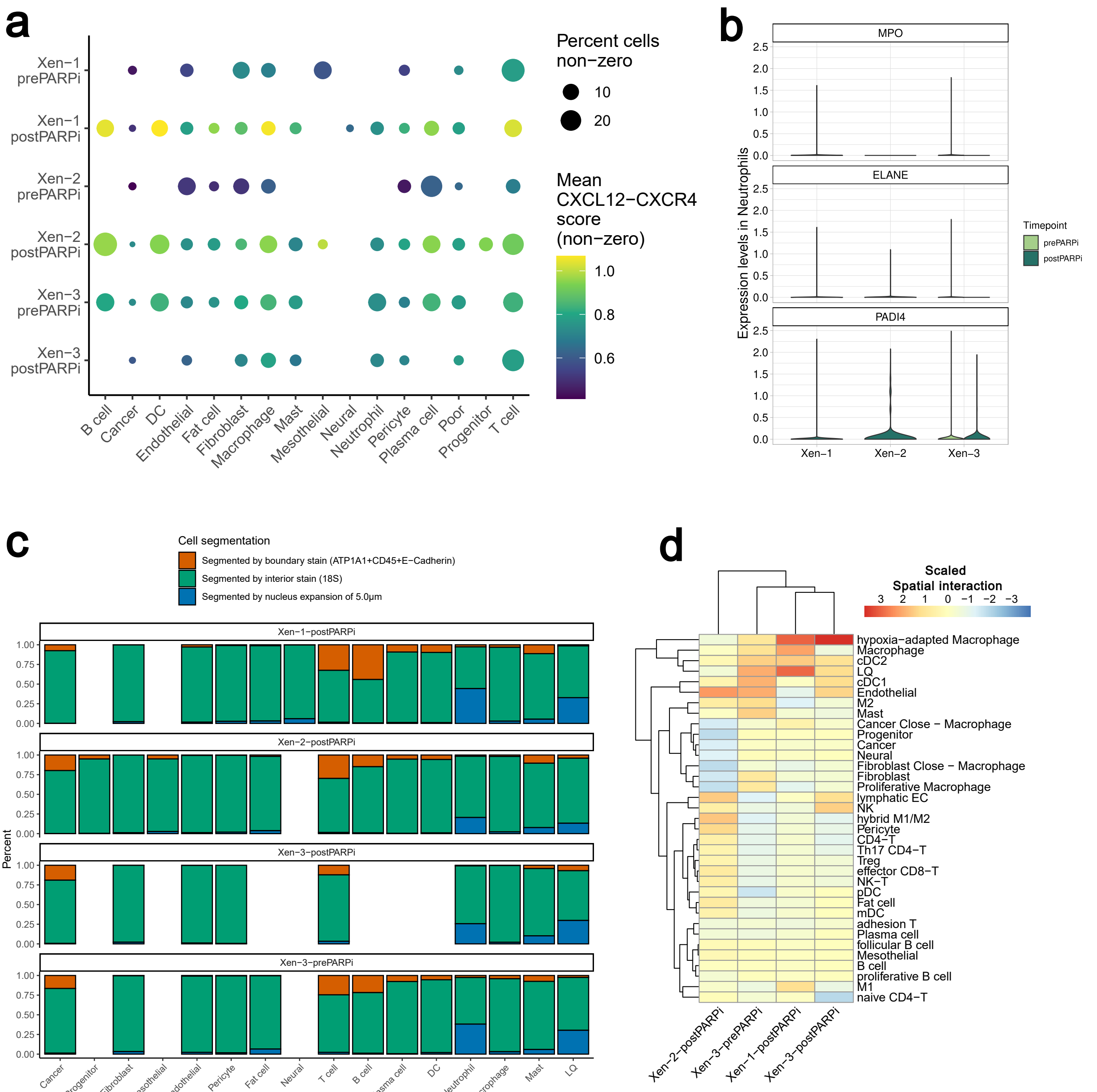

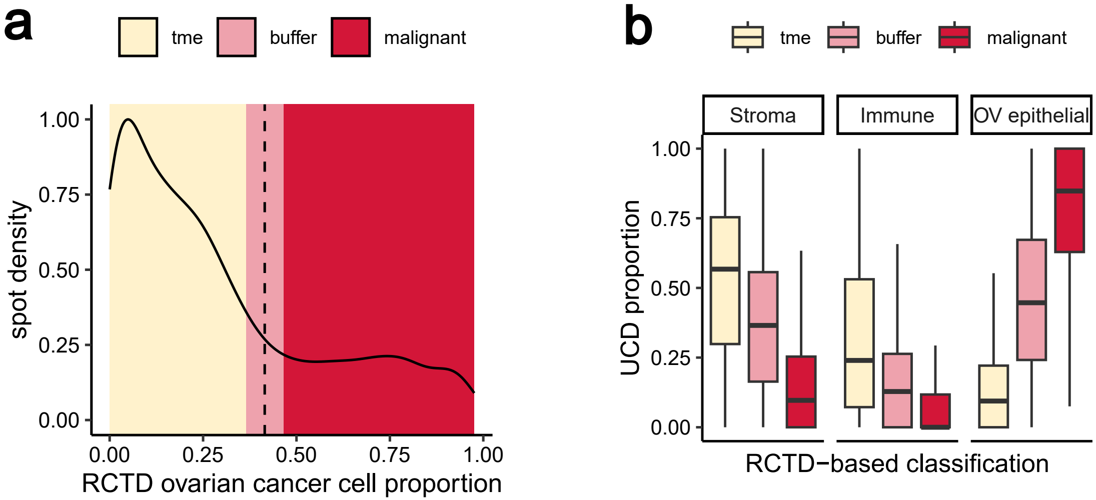

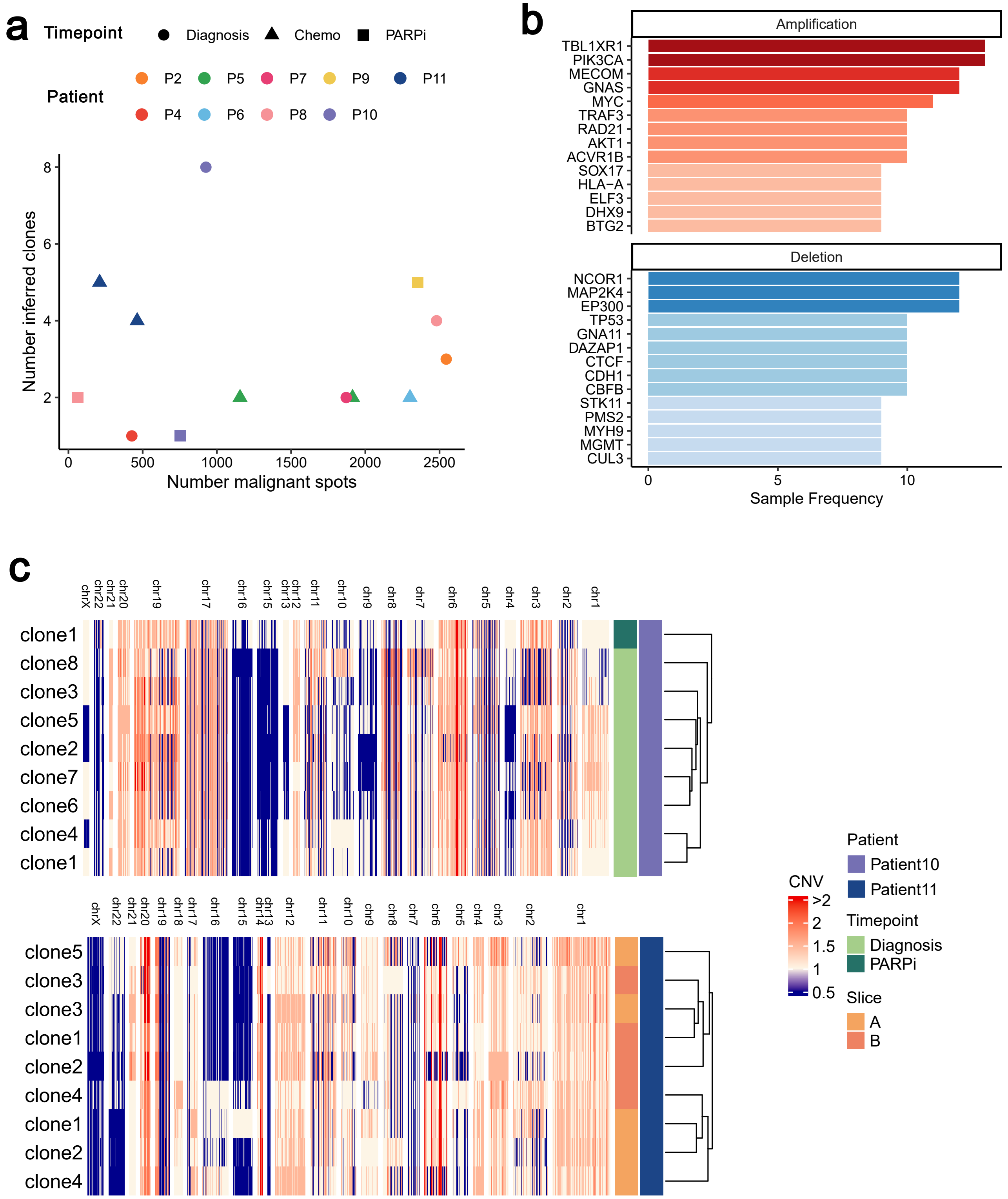
